# A patatin-like phospholipase mediates *Rickettsia parkeri* escape from host membranes

**DOI:** 10.1101/2021.10.21.465009

**Authors:** Gina M. Borgo, Thomas P. Burke, Cuong J. Tran, Nicholas T. N. Lo, Patrik Engström, Matthew D. Welch

## Abstract

Spotted fever group *Rickettsia* species are arthropod-borne obligate intracellular bacteria that can cause mild to severe human disease. These bacteria invade host cells, replicate in the cell cytosol, and then spread from cell to cell. To access the host cytosol and avoid detection by immune surveillance mechanisms, these pathogens must have evolved efficient ways to escape membrane-bound vacuoles. Although *Rickettsia* are predicted to express factors that disrupt host membranes, little is known about how and when these proteins function during infection. Here, we investigated the role of a *Rickettsia* patatin-like phospholipase A2 enzyme (Pat1) during host cell infection by characterizing a *Rickettsia parkeri* mutant with a transposon insertion in the *pat1* gene. We show that Pat1 is important for infection in a mouse model and in host cells. We further show that Pat1 is critical for efficiently escaping from the single and double membrane-bound vacuoles into the host cytosol, and for avoiding host galectins that mark damaged membranes. In the host cytosol, Pat1 is important for avoiding host polyubiquitin, preventing recruitment of autophagy receptor p62, and promoting actin-based motility and cell-cell spread. Our results show that Pat1 plays critical roles in escaping host membranes and promoting cell-cell spread during *R. parkeri* infection and suggest diverse roles for patatin-like phospholipases in facilitating microbial infection.

**Importance:** Spotted fever group *Rickettsia* are bacteria that reside in ticks and can be transmitted to mammalian hosts, including humans. Severe disease is characterized by high fever, headache, and rash, and results in occasional mortality despite available treatment. *Rickettsia* interact with host cell membranes while invading cells, escaping into the cytosol, and evading cellular defenses. Bacterial phospholipase enzymes have been proposed as critical factors for targeting host cell membranes, however the specific roles of rickettsial phospholipases are not well defined. We investigated the contribution of one conserved patatin-like phospholipase, Pat1, in *Rickettsia parkeri*. We observed that Pat1 is important for virulence in an animal model. Moreover, Pat1 plays critical roles in host cells by facilitating access to the cell cytosol, inhibiting detection by host defense pathways, and promoting cell-cell spread. Our study indicates that Pat1 performs several critical functions, suggesting a broad role for phospholipases throughout the *Rickettsia* lifecycle.

## Introduction

Spotted fever group (SFG) *Rickettsia* species are Gram-negative, obligate intracellular bacteria that infect tick vectors and can be transmitted by tick bites to vertebrate hosts (1). SFG *Rickettsia* that can cause disease in humans include *R. rickettsii*, the causative agent of Rocky Mountain spotted fever, a disease characterized by high fever, neurological symptoms, organ failure, and possible fatality if left untreated (2, 3). Disease-causing SFG *Rickettsia* also include species such as *R. parkeri*, which causes milder eschar-associated rickettsiosis characterized by fever and a skin lesion (eschar) at the site of the tick bite, but is not documented to cause fatality (2, 4, 5). Because *R. parkeri* can be studied under biosafety level 2 (BSL2) conditions, it is emerging as a model for understanding the molecular determinants of SFG *Rickettsia* pathogenicity.

*R. parkeri* targets macrophages (4–8) as well as endothelial cells (6, 7, 9) during infection in humans and animal models. Upon invasion of host cells, bacteria escape from the primary vacuole into the cytosol, where they replicate (10, 11). Bacteria then initiate actin-based motility and move to the plasma membrane, where they enter into protrusions that are engulfed into neighboring cells (12). This necessitates another escape event from a double-membrane secondary vacuole into the cytosol, completing the intracellular life cycle (10, 11). Other bacteria with a similar life cycle utilize pore-forming proteins and phospholipases to escape from the primary and/or secondary vacuole. For example, *Shigella flexneri* uses the IpaB-IpaC translocon to form pores that facilitate membrane rupture (13–18). *Listeria monocytogenes* utilizes the cholesterol-dependent cytolysin listeriolysin O (LLO) (19–22) and two phospholipase C enzymes, PlcA and PlcB, to escape from primary and secondary vacuoles (19, 23–27). It is likely that *Rickettsia* also utilizes at least one protein that can directly disrupt the vacuolar membrane to mediate escape.

SFG *Rickettsia* genomes encode two types of phospholipase enzymes, phospholipase D (PLD) and up to two patatin-like phospholipase A2 (PLA2) enzymes (Pat1 and Pat2) (28, 29). PLD is dispensable for vacuolar escape, as a *pld* mutant in *R. prowazekii* showed no delay in escape (30), even though exogenous PLD expression in *Salmonella enterica* was sufficient to facilitate escape (31). In contrast, evidence suggests a possible role for PLA2 enzymes in escape. For example, PLA2 activity from *R. prowazekii* was demonstrated to target host phospholipids throughout infection (32, 33). Furthermore, pretreatment of bacteria with either a PLA2 inhibitor, or anti-Pat1 or anti-Pat2 antibodies, reduced plaque number for both *R. rickettsii* (34–36) and *R. typhi* (28, 37), and increased colocalization of *R. typhi* with the lysosomal marker LAMP-1 (37). This suggests that Pat1 and Pat2 are important for infection and avoidance of trafficking to the lysosome. Nevertheless, the precise role of PLA2 enzymes in rickettsial vacuolar escape has remained unclear.

Phospholipase activity and escape from the vacuole may also be important to enable downstream life cycle events, such as actin-based motility, which requires access to actin in the host cell cytosol. Another is avoidance of anti-bacterial autophagy (also called xenophagy). Autophagy can be initiated via polyubiquitination of cytosolic bacteria (38–40) and subsequent recruitment of autophagy receptors (41) such as p62/Sequestome 1 (SQSTM1) (hereafter referred to as p62) (42–44) and NDP52 (nuclear dot protein 52 (NDP52)/calcium-binding and coiled-coil domain 2 (CALCOCO2)) (hereafter referred to as NDP52) (42, 45, 46). Autophagy receptors recognize polyubiquitinated bacteria and interact with microtubule-associated protein 1A/1B-light chain 3 (LC3), which marks nascent and mature autophagosomal membranes that can enclose bacteria and deliver them to the lysosome (38, 47, 48). Bacterial phospholipases may facilitate autophagy avoidance through manipulation of phospholipids needed for autophagosome formation, such as with *L. monocytogenes* PlcA targeting of phosphatidylinositol 3-phosphate (PI(3)P) to block LC3 lipidation (49, 50).

Autophagy can also be initiated by membrane damage to the bacteria-containing vacuole, which exposes glycans internalized from the host cell surface that are recognized by host cytosolic glycan-binding galectin (Gal) proteins (51). Gal3 and Gal8 can target damaged vacuolar compartments during infection with *L. monocytogenes* (52, 53) and *S. flexneri* (52, 54, 55), as well as during infection with bacteria that typically reside in membrane-bound compartments such as *Legionella pneumophila* (56), *S. enterica* (52, 55), *Coxiella burnetti* (57), and *Mycobacterium tuberculosis* (58). Importantly, membrane remnants marked by Gal3 or Gal8 colocalize with polyubiquitin (54, 58, 59), autophagy receptors p62 (54, 58) and NDP52 (55, 57), and LC3 (54, 55, 57, 58). Nevertheless, it remains unknown if rickettsial phospholipases are important to evade autophagy.

To better understand the role of PLA2 enzymes during SFG *Rickettsia* infection, we characterized a *R. parkeri* mutant with a transposon insertion in the single PLA2-encoding gene *pat1*. We found that Pat1 is critical throughout infection for escaping host membranes, avoiding targeting by autophagy, and spreading to neighboring cells. These results suggest that Pat1 is a key bacterial factor involved in interacting with host membranes and avoiding detection in host cells.

## Results

### Pat1 is important for infection of host cells and contributes to virulence in mice

To determine the role of Pat1 during infection, we used a *R. parkeri* mutant with a transposon insertion in the *pat1* gene (*pat1*::Tn) that was previously isolated in a screen for mutants with reduced plaque size (60). We complemented the *pat1*::Tn mutation by generating a strain (*pat1*::Tn *pat1^+^*) that contains a second transposon encoding full length *pat1* plus the intergenic regions immediately 5’ and 3’ to the gene (predicted to contain the native promoter and terminator) (**Fig. 1A**). Using an antibody that recognizes *R. parkeri* Pat1 by western blotting, we observed a band at the predicted molecular weight of 55 kD for Pat1 in WT bacteria, no corresponding band in the *pat1*::Tn mutant, and a restoration of the band in the *pat1*::Tn *pat1^+^*-complemented mutant (**Fig. 1B**). The absence of detectable Pat1 protein in the mutant suggests it is a null mutant. Because the *pat1*::Tn mutant was initially identified based on its small-plaque phenotype, we next compared plaque area for WT, mutant, and complemented mutant strains. In comparison with WT, the *pat1*::Tn mutant showed significantly smaller plaques, and plaque area was rescued in the complemented mutant (**Fig. 1C, D**). This demonstrates that the observed reduction in plaque area was caused by loss of *pat1*.

**Fig. 1.**
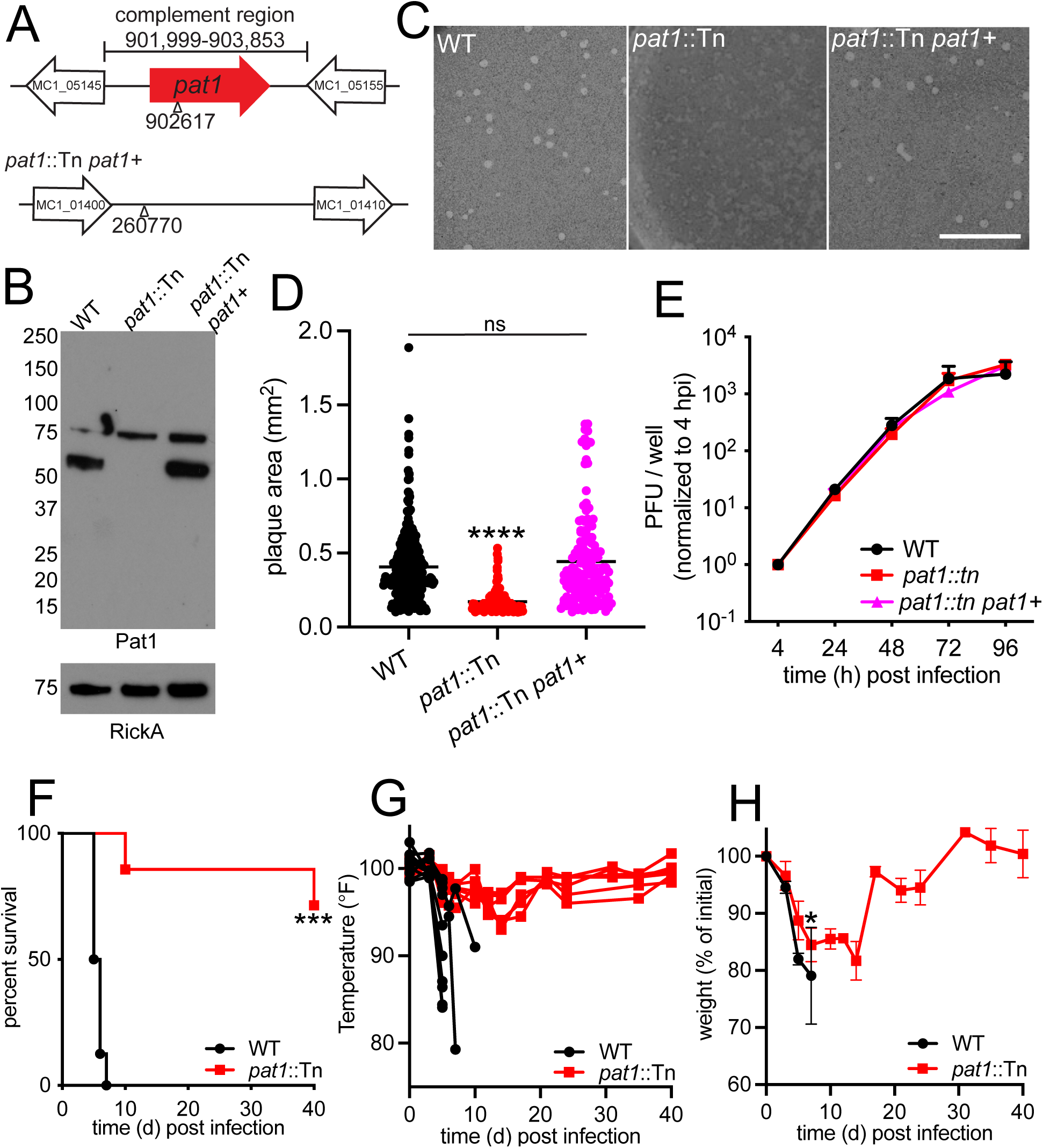
Pat1 is important for infection of cells and in mice. (A) Genomic loci of *pat1* (top) and *pat1* insertion site for complementation (bottom). Triangle represents transposon insertion sites, and nucleotide numbers indicate the position in the *R. parkeri* genome. Genes upstream and downstream are included to show intergenic regions. (B) Western blot of purified *R. parkeri* strains, WT, *pat1*::Tn, and complemented strain (*pat1*::Tn *pat1^+^*), probed with anti-Pat1 antibody; probing with anti-RickA was used as a loading control. Pat1 has a predicted size of 55kD. Numbers on left are molecular weight in kD. (C) Images of plaques stained with neutral red at 6 dpi. Scale bar 10 mm. (D) Plaque areas in Vero cells infected with WT, *pat1*::Tn, and complemented strain (n=2 independent experiments; 50-80 plaques per experiment). (E) Growth curve of WT, *pat1*::Tn, and complemented strain in HMECs (n=3 independent experiments). (F) Survival of *Ifnar1^-/-^*;*Ifngr1^-/-^* mice infected intravenously (i.v.) with 5×10^6^ WT or *pat1*::Tn mutant (n=8 mice for WT, n=7 mice for *pat1*::Tn, data represents 2 independent experiments). (G) Temperature changes over time in i.v. infection of *Ifnar1^-/-^*;*Ifngr1^-/-^* mice with 5×10^6^ WT or *pat1*::Tn mutant bacteria; graphs represent data from individual mice. (H) Weight change over time expressed as percent change from initial weight in i.v. infection of *Ifnar1^-/-^*;*Ifngr1^-/-^* mice with 5×10^6^ WT or *pat1*::Tn mutant bacteria. Data in (D) and (E) are mean ± SEM; ****p<0.0001 relative to WT (one-way ANOVA), ns=not significant. Data in (F) were analyzed using a log-rank (Mantel-Cox) test ***p<0.001. Data in (H) were analyzed using a two-way ANOVA from 0 to 7 dpi.

To further determine if Pat1 plays a role in bacterial replication, growth curves measuring PFU were performed in human microvascular endothelial cells (HMECs). There were no differences in bacterial replication kinetics for WT, *pat1*::Tn and *pat1*::Tn *pat1^+^* -complemented strains in HMECs (**Fig. 1E**). These data indicate that the transposon disruption of *pat1* does not interfere with intracellular growth in these cell lines.

We next examined the contribution of Pat1 to virulence *in vivo* using mice lacking the receptors for both IFN-I (*Ifnar1*) or IFN-*γ* (*Ifngr1*) (*Ifnar1^-/-^*;*Ifngr1^-/-^* double knock out mice), which succumb to infection with WT *R. parkeri* and can be used to investigate the importance of bacterial genes to virulence (8, 61). Mice infected intravenously (i.v.) with 5×10^6^ PFU WT bacteria showed a rapid drop in temperature and body weight following infection and did not survive past day 8 (**Fig 1F, G, H**). In contrast, mice infected i.v. with the *pat1*::Tn mutant maintained a steady temperature following infection, showed an initial drop in weight that stabilized around 2 weeks post infection before increasing, and the majority survived until the end of the experiment (day 40) (**Fig. 1F, G, H**). These results indicate that Pat1 is an important virulence factor.

### Pat1 promotes efficient escape from the vacuole post-invasion

Because *R. typhi* Pat1 and Pat2 had previously been implicated in avoidance of trafficking to lysosomes (37), we sought to determine if the *R. parkeri pat1*::Tn mutant was impaired in its ability to escape from the primary vacuole during infection. To evaluate the role of Pat1 in vacuolar escape, we used transmission electron microscopy (TEM) to investigate whether host membranes surrounded intracellular bacteria 1 h post infection (hpi) in HMECs. This time point was chosen because prior studies reported escape from the vacuole by 30 min post infection (mpi) for *R. typhi* (37), *R. prowazekii* (30), and *R conorii* (*62*). At this timepoint, significantly more WT bacteria were found free in the cytosol (74%) compared with the *pat1*::Tn mutant (38%) (**Fig. 2A, B)**. Moreover, significantly fewer WT bacteria were found within membranes (25% in single membranes, 1% in double membranes) in comparison with the *pat1*::Tn mutant (50% in single membranes, 12% within double membranes). Differences in vacuolar localization were not due to differences in invasion of host cells, as both WT and the *pat1*::Tn mutant entered cells with similar kinetics (**Fig. 2C**). These results suggest that Pat1 facilitates escape from membranes following invasion.

**Fig. 2.**
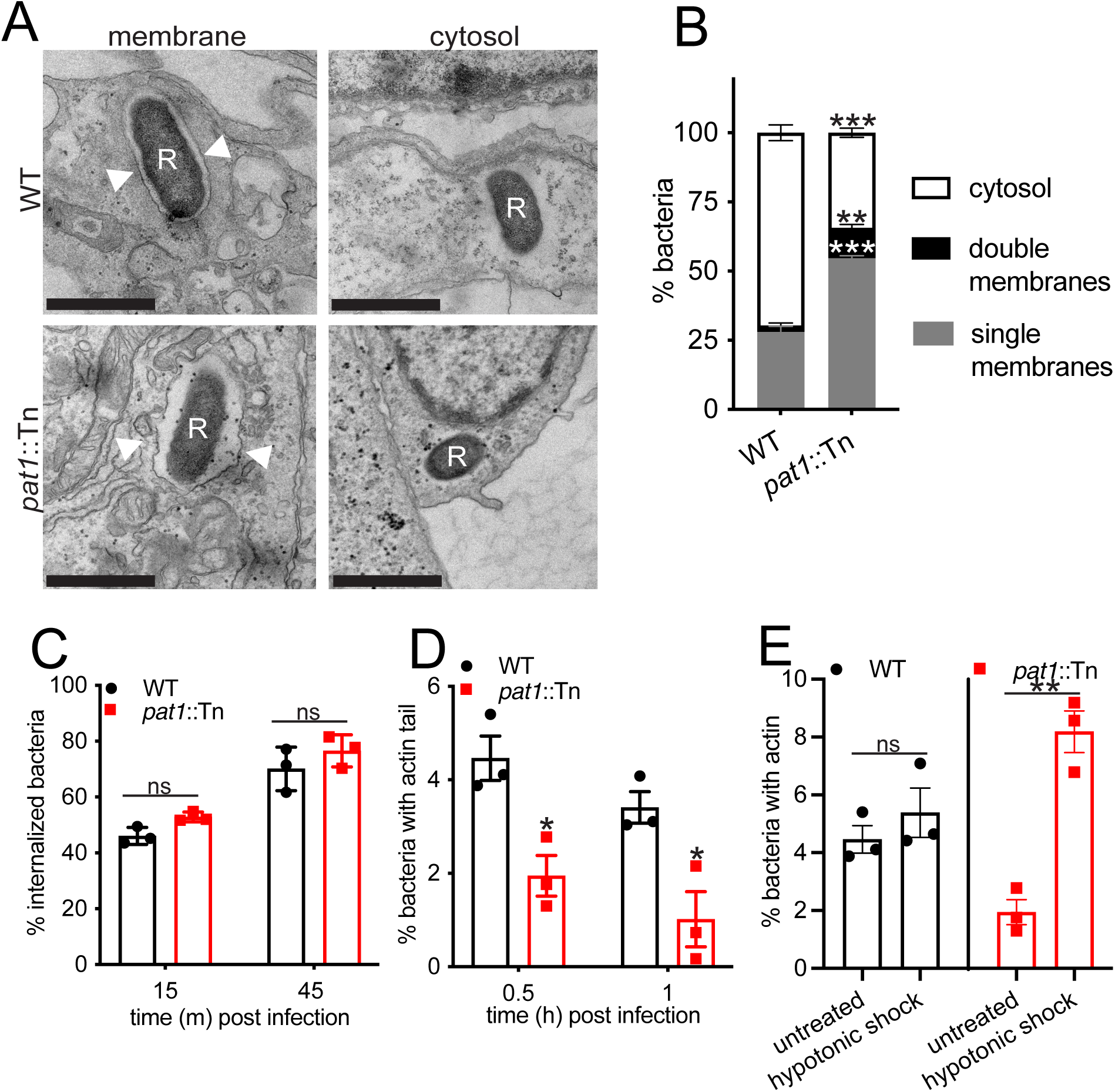
Pat1 facilitates escape from single and double membrane compartments following invasion. (A) TEM images of WT and *pat1*::Tn mutant bacteria in HMECs at 1 hpi. “R” indicates *R. parkeri* and arrowheads point to membrane surrounding the bacteria. Scale bar 1 µm. (B) Quantification of (A), percentage of single and double membrane-bound or cytosolic bacteria (WT=80 bacteria, *pat1*::Tn=88 bacteria, n=3 independent experiments). (C) Percent of bacteria internalized at 15 mpi and 45 mpi (images not shown). (D) Percent of bacteria with actin tails at 30 mpi and 1 hpi (images not shown). (E) Percent of bacteria with actin in untreated cells or cells that have undergone hypotonic shock treatment to lyse vacuoles (images not shown). All data represents n=3 independent experiments. Data in (B, C, D) are mean ± SEM; ***p<0.001 **p<0.01 *p<0.05, ns=not significant relative to WT (unpaired t-test). Data in (E) are mean ± SEM; *p<0.01 relative to untreated (paired t-test).

We further hypothesized that the increased localization of the *pat1*::Tn mutant within membranes impaired access to the cytosol, particularly to the pool of actin, interfering with actin-based motility. To test this hypothesis, we quantified the number of bacteria with actin tails at 30 mpi and 1 hpi. Approximately 3-4% of WT bacteria were associated with actin tails, in keeping with previous reports (63, 64). The frequency of *pat1*::Tn mutant association with actin tails was half that of WT at both time points (**Fig. 2D**). These results suggest that failure of the *pat1*::Tn mutant to escape from the vacuole leads to reduction in actin-based motility. To confirm that the reduced frequency of actin-based motility resulted from bacteria being trapped within membranes, we used hypotonic shock (alternating treatment with hypertonic and then hypotonic solutions) to lyse primary vacuoles (65, 66) and more efficiently deliver bacteria to the cytosol. When cells infected with WT bacteria were subjected to hypotonic shock at 5 mpi, there was no significant increase in the percentage of bacteria with actin tails at 30 mpi, suggesting that WT bacteria optimally access the cytosol following invasion (**Fig. 2E**). In contrast, hypotonic shock significantly increased the percentage of *pat1*::Tn mutant bacteria with actin tails. These results confirm that reduced frequency of actin-based motility in the *pat1::tn* mutant is due to entrapment in the primary vacuole.

### Pat1 contributes to autophagy avoidance

The presence of a marked fraction (12%) of *pat1*::Tn mutant bacteria in double-membrane compartments at 1 hpi could not be explained by failure to escape from the vacuole, suggesting the possibility that bacteria were targeted by host cell autophagy. Because an initial step of anti-bacterial autophagy is recognition and ubiquitylation of the bacterial surface (40), we first tested for bacterial association with polyubiquitin in infected HMECs at 0-2 hpi. Whereas fewer than 2% of WT bacteria were polyubiquitin-positive from 0-2 hpi, the percentage of polyubiquitin-positive *pat1*::Tn mutant bacteria was significantly higher and increased (from about 6% at 0 hpi to about 16% at 1 hpi), before falling slightly (**Fig. 3A, B)**. Complementation of the *pat1*::Tn mutant reduced the percent of polyubiquitin-positive bacteria to levels seen with WT (**Fig. 3C**). This suggests that Pat1 reduces recognition of bacteria by the host ubiquitylation machinery.

**Fig. 3.**
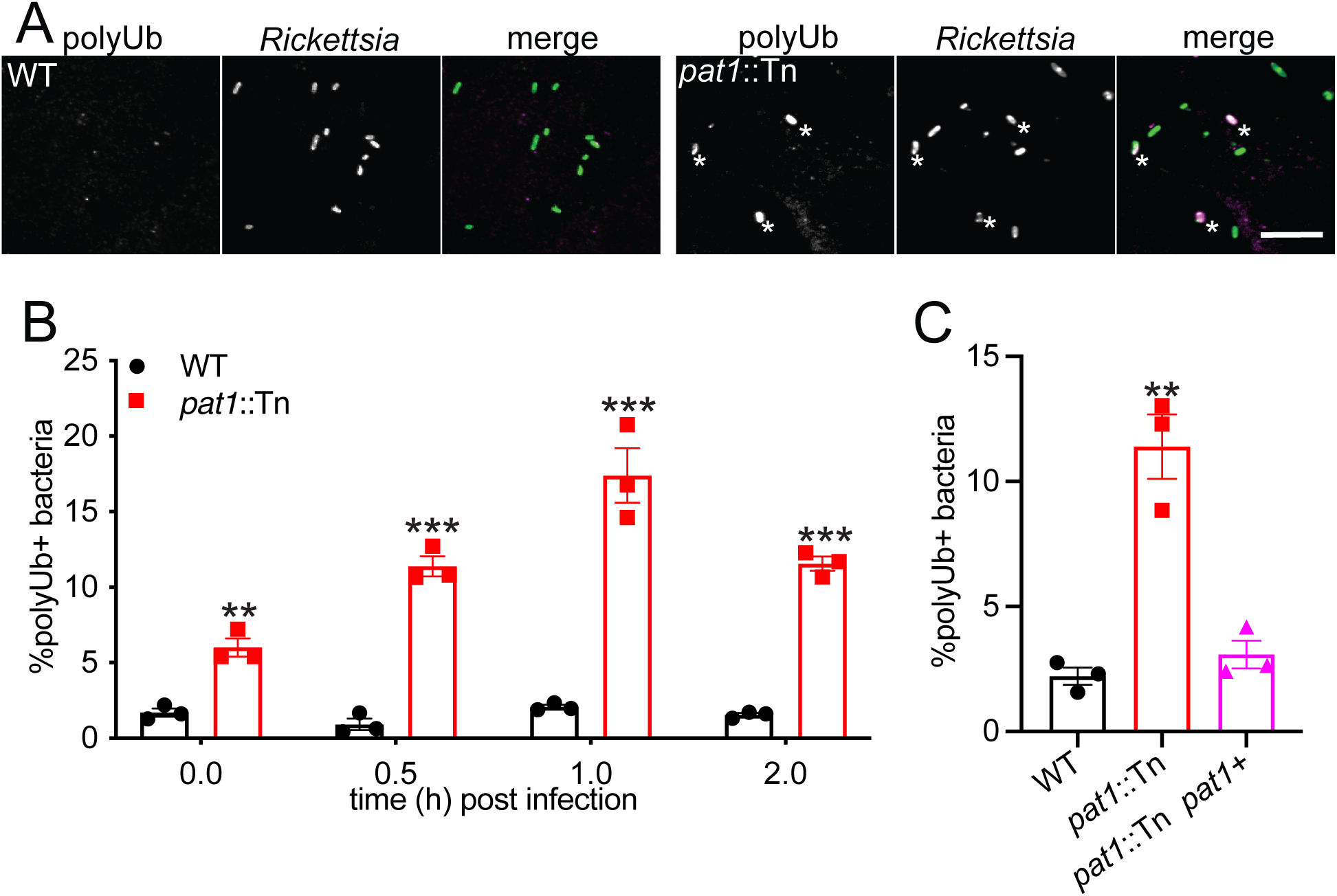
Pat1 contributes to avoidance of polyubiquitin recruitment. (A) Images of polyubiquitin (polyUb; magenta in merge) in HMECs infected with WT and *pat1*::Tn bacteria (green in merge) at 1 hpi. Asterisk denotes colocalization between bacterium and polyUb. Scale bar is 5 µm. (B) Quantification of (A), percentage of polyUb-positive bacteria at the indicated time points. (C) Percent of polyUb-positive bacteria in HMECs infected with WT, *pat1*::Tn, or complemented mutant (*pat1*::Tn *pat1+*) at 1 hpi. All data represent n=3 independent experiments. Data in (B) are mean ± SEM; ***p<0.001 **p<0.01*p<0.05 relative to WT (unpaired t-test). Data in (C) are mean ± SEM; ***p<0.001 **p<0.01 *p<0.05 relative to WT (one-way ANOVA).

To further examine whether bacteria were targeted by the autophagy machinery, we examined the recruitment of autophagy receptors p62 and NDP52, as well as LC3 at 1 hpi, the time point with the most polyubiquitin-positive bacteria. Compared with WT (fewer than 2% stained with these markers), markedly more of the *pat1*::Tn mutant were positive for p62 (10%) and NDP52 (6%) (**Fig. 4A, B, C**). Moreover, more of the *pat1*::Tn mutant bacteria colocalized with LC3 at 1 and 2 hpi (**Fig. 4D, E**). Interestingly, the increased recruitment of LC3 to the *pat1::tn* mutant preceded increased colocalization of the mutant with LAMP-1, a marker for late endosomal and lysosomal compartments (**Fig. S1A, B**). These results suggest that Pat1 is important for counteracting the recruitment of autophagy receptors and targeting to autophagosomes and lysosomes.

**Fig. 4.**
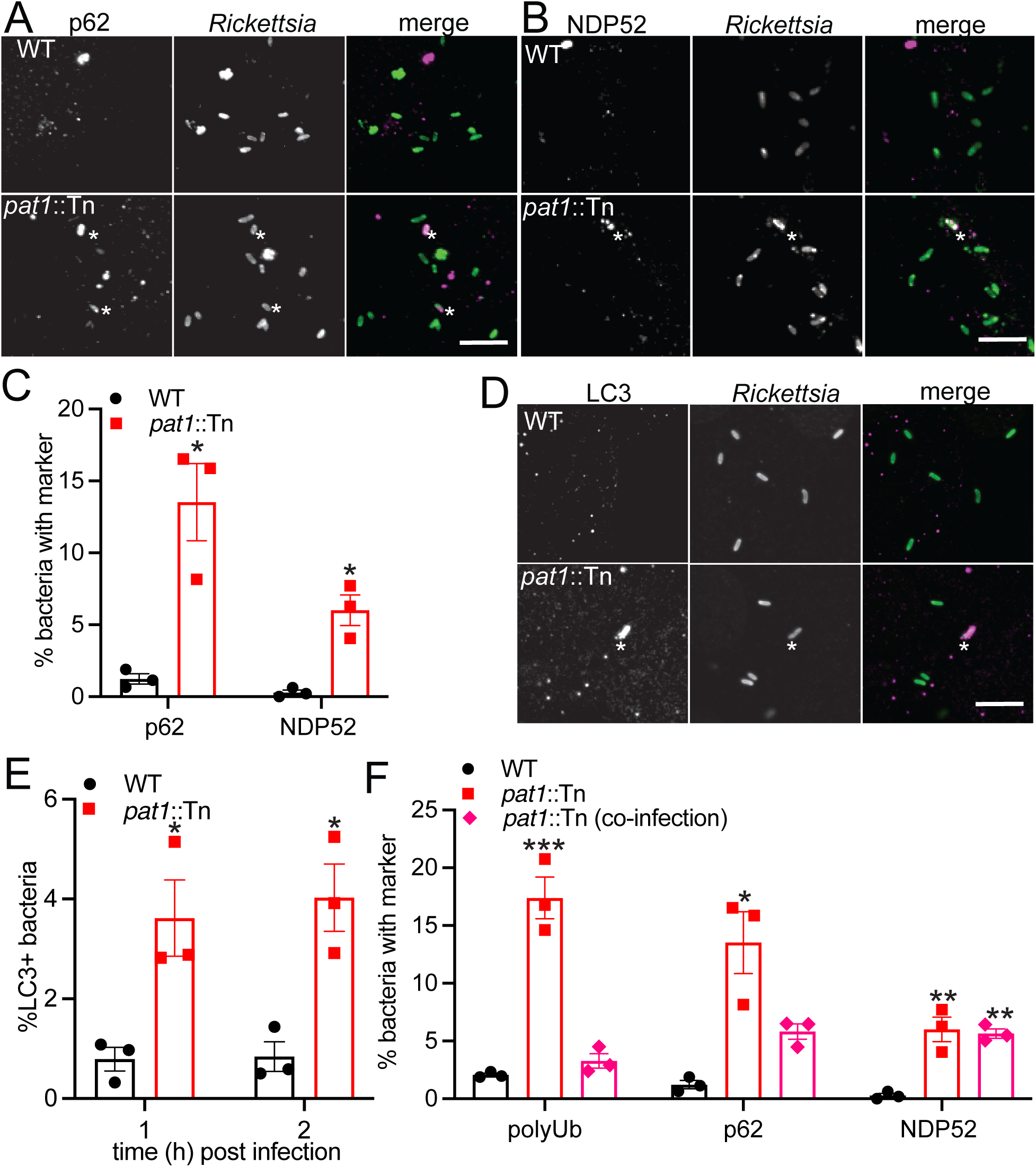
Pat1 enables evasion of recognition by autophagy. Images of autophagy receptors (A) NDP52 (left; magenta in merge) and (B) p62 (right; magenta in merge) in WT and *pat1*::Tn (green in merge) infected HMECs. Asterisk denotes colocalization between bacterium and p62 or NDP52. (C) Quantification of (A) and (B), percentage of bacteria staining for NDP52 or p62 at 1 hpi. (D) Images of LC3 (magenta) in HMECs infected with WT and *pat1*::Tn bacteria (green) at 1 hpi. Asterisk denotes colocalization between bacterium and LC3. (E) Quantification of percent bacteria staining for LC3 at 1 hpi (images in (D)) and 2 hpi (images not shown). (F) Percentage colocalization of bacteria with polyUb, NDP52, and p62 in HMECs infected with WT, *pat1*::Tn mutant, or co-infected with WT and *pat1*::Tn mutant. For co-infections, quantification is for *pat1*::Tn bacteria only. All data represents n=3 independent experiments. Data in (C, E) are mean ± SEM; ***p<0.001 **p<0.01*p<0.05 relative to WT (unpaired t-test). Data in (F) are mean ± SEM; ***p<0.001 **p<0.01 *p<0.05 relative to WT (one-way ANOVA). Scale bars in (A, B, D) are 5 µm.

Because previous studies suggested that Pat1 is secreted into the host cell (37), we also sought to further ascertain whether Pat1 was counteracting ubiquitylation and targeting by the autophagy machinery by acting locally on the bacterium producing the protein, and/or by acting at a distance on other bacteria. To test this, we co-infected HMECs with WT bacteria expressing 2xTagBFP and with *pat1*::Tn mutant bacteria, and quantified colocalization of *pat1*::Tn bacteria with polyubiquitin, NDP52, and p62. The *pat1*::Tn mutant exhibited significantly reduced colocalization with polyubiquitin and p62 (but not NDP52) in co-infected cells compared with cells infected with the *pat1*::Tn mutant only (**Fig. 4F**). These results suggest that Pat1 is secreted and can function at a distance to reduce bacterial targeting with polyubiquitin and p62.

### Pat1 antagonizes bacterial association with damaged membranes that recruit Gal3 and NDP52

It remained unclear whether polyubiquitin and the autophagy machinery were associated with bacteria enclosed in damaged vacuolar membranes or those free in the cytosol. To determine whether polyubiquitin, NDP52, and p62 were present at damaged vacuoles at 1 hpi, we quantified the percentage of bacteria staining for polyubiquitin, NDP52, or p62, in cells transiently expressing Gal3-mCherry or Gal8-mCherry to mark damaged membranes (52, 55) (**Fig. 5A, B)**. A small fraction (0.5%) of WT bacteria colocalized with Gal3-mCherry or Gal8-mCherry, whereas significantly more *pat1::tn* mutant bacteria colocalized with Gal3-mCherry or Gal8-mCherry (∼2.5%) (**Fig. 5C, D**). A significantly higher fraction of the *pat1*::Tn mutant bacteria that stained for NDP52 also colocalized with either Gal protein (∼50%) (**Fig. 5E**). Some *pat1*::Tn mutant bacteria that stained for p62 also colocalized with Gal3-mCherry (∼5%) or Gal8-mCherry (∼10%) (**Fig. 5E**). We rarely observed colocalization of the *pat1*::Tn mutant bacteria with both polyubiquitin and either Gal protein (**Fig. 5E**). Interestingly, although the *pat1*::Tn mutant was more frequently associated with Gal3-mCherry or Gal8-mCherry, in *pat1*::Tn mutant-infected cells we observed fewer clusters of Gal3-mCherry (**Fig. S2A, C)** or Gal8-mCherry (**Fig. S2B, D)** not associated with bacteria, consistent with reduced overall membrane damage compared with cells infected with WT bacteria.

**Fig. 5.**
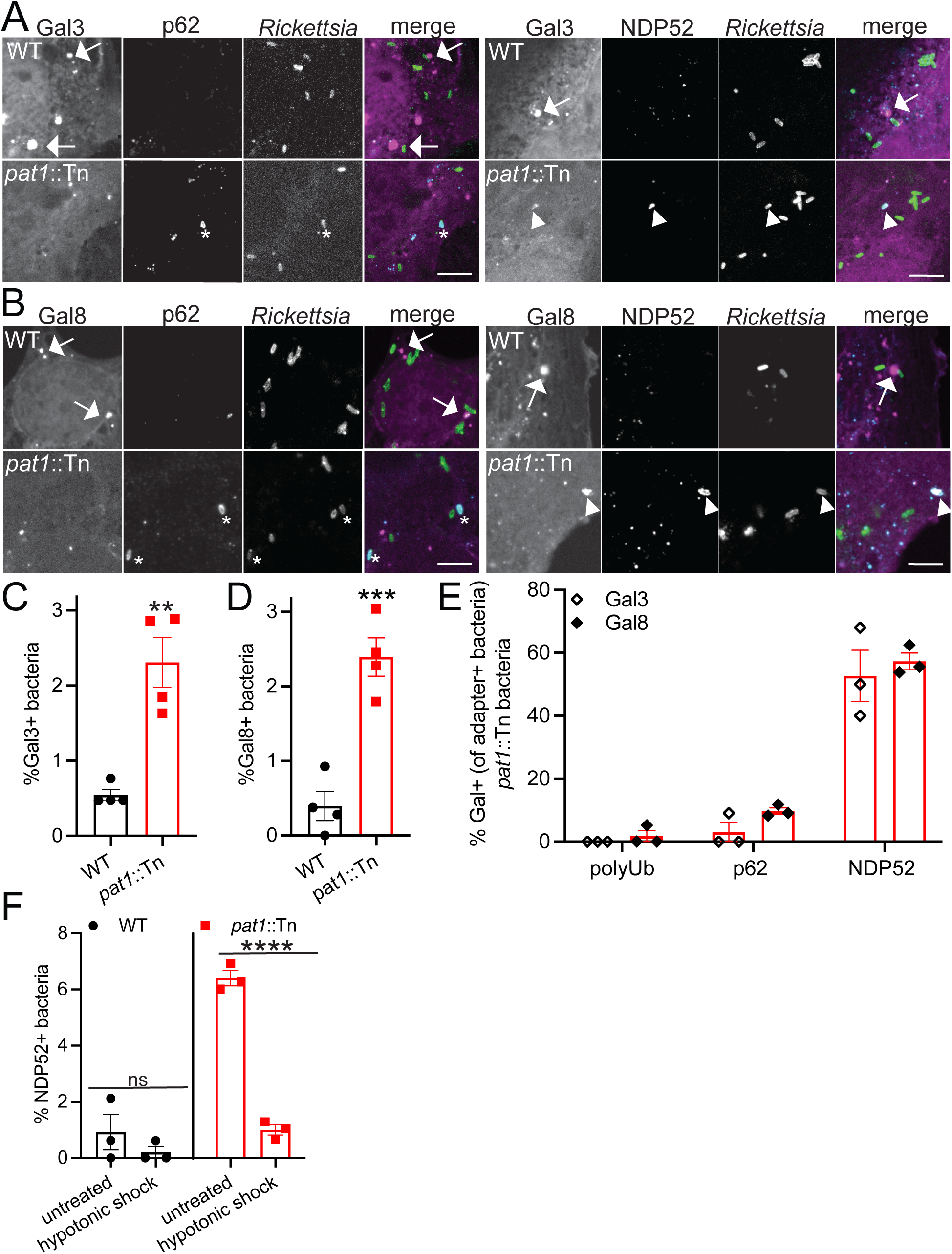
Pat1 is important for avoiding bacterial association with damaged membranes. (A) Images of Gal3-mCherry (magenta in merge) and autophagy receptors p62 (left; cyan in merge) and NDP52 (right; cyan in merge) in HMECs infected with WT or *pat1*::Tn bacteria (green in merge) at 1 hpi. Arrows indicate large Gal3 positive clusters near bacteria, asterisk indicates bacteria that are p62 positive and Gal3 negative, arrowheads indicate colocalization between NDP52, Gal3, and bacteria. (B) Images of Gal8-mCherry (magenta in merge) and autophagy receptors p62 (left; cyan in merge) and NDP52 (right; cyan in merge) in HMECs infected with WT or *pat1*::Tn bacteria (green in merge) at 1 hpi. Arrows indicate large Gal8 positive clusters near bacteria, asterisk indicates bacteria that are p62 positive and Gal8 negative, arrowheads indicate colocalization between NDP52, Gal8, and bacteria. Scale bar for (A) is 5 µm and (B) is 3 µm. (C) Quantification of (A), percentage of bacteria positive for Gal3 (n=4). (D) Quantification of (B), percentage of bacteria positive for Gal8 (n=4). (E) Quantification of (A) and (B), percent of polyUb, p62, or NDP52 positive *pat1*::Tn bacteria that are also positive for Gal3 (open diamond) or Gal8 (filled diamond) (n=3 for all markers). (F) Percent of bacteria positive for NDP52 in untreated cells or cells that undergo hypotonic lysis of vesicles (n=3). Data in (C) and (D) represent n=4 independent experiments and data in (E) and (F) represent n=3 independent experiments. Data in (C), (D) are mean ± SEM; ***p<0.001 **p<0.01 relative to WT (unpaired t-test). Data in (F) are mean ± SEM; **p<0.01 relative to untreated (paired t-test).

To test whether NDP52 associated with bacteria or damaged membranes, we performed the hypotonic shock treatment (**Fig. 2E**) to release bacteria from host vacuoles. Cells infected with WT and *pat1*::Tn bacteria were subjected to hypotonic shock treatment at 5 mpi, and then at 30 mpi, we quantified the number of bacteria that colocalized with NDP52. Hypotonic shock significantly reduced the percent colocalization of the *pat1*::Tn with NDP52 (from ∼6% in untreated cells to ∼1% in treated cells), while fewer than 1% of WT bacteria colocalized with NDP52 regardless of treatment (**Fig. 5F**). Together, these results support the conclusion that Pat1 promotes efficient escape from vacuolar membranes and enables avoidance of targeting by Gal3, Gal8, and NDP52.

### Pat1 facilitates actin-based motility and spread into neighboring cells late in infection

Although Pat1 is dispensable for bacterial replication kinetics, it promotes plaque formation, suggesting that Pat1 may function in cell-cell spread. To initially assess if Pat1 is important for spread, we used an infectious focus assay, in which the number of infected host cells per focus of infection was quantified at 28 hpi to measure spread efficiency (67). Compared with WT bacteria (∼4.5 cells per focus), the *pat1*::Tn mutant infected significantly fewer cells (∼3.5 cells per focus) (**Fig. 6A, B**). This suggested that Pat1 is important for spread. To further assess cell-cell spread, we carried out a “mixed cell” assay (67) in which “primary” A549 cells stably expressing the TagRFP-T-farnesyl plasma-membrane marker (A549-TRTF) were infected for 1 h, detached from the plate, and mixed with unlabeled “secondary” A549 cells (**Fig. 6C, D).** The percent of bacteria in the primary A549-TRTF cell and secondary cell were quantified at 32 hpi. We observed that 50% of WT bacteria were found in primary cells and 50% had spread into secondary cells. In contrast, ∼85% of *pat1*::Tn mutant bacteria remained in primary cells and only ∼15% had spread to secondary cells (**Fig. 6C, E**). This confirms that Pat1 is important for cell-cell spread.

**Fig. 6.**
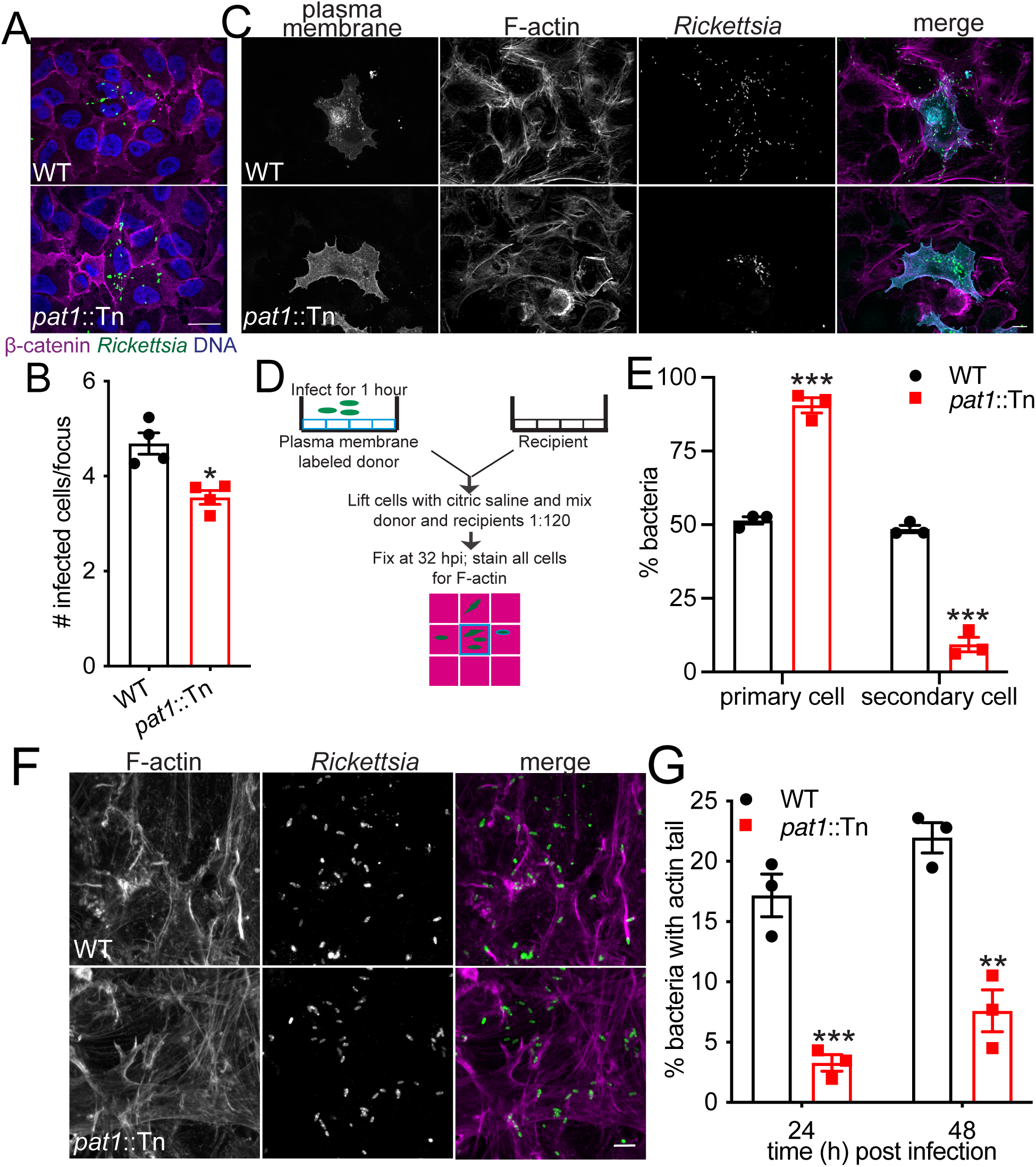
Pat1 is important for cell-cell spread and facilitates actin-based motility. (A) Images of infectious foci formed by WT or *pat1*::Tn mutant in A549 cells at 28 hpi (magenta, β-catenin; green, bacteria; blue, nuclei). Scale bar is 10 μm. (B) Quantification of (A), number of infected cells per focus (n=4). (C) Images of mixed cell assay depicted in (D) showing plasma membrane (A549-TRTF; cyan in merge), F-actin (magenta in merge), and bacteria (green in merge), adapted from (67). Scale bar is 10 µm. (E) Percent bacteria in primary and secondary cells quantified from (C) (n=3). (F) Images of actin tails (F-actin; magenta in merge) and bacteria (green in merge) in HMECs at 24 hpi. (G) Percent of bacteria with actin tails at 24 hpi (from (F) and 48 hpi in HMECs (images not shown) (n=3). Scale bar is 5 µm. Data in (B) represents n=4 independent experiments and data in (E) and (G) represent n=3 independent experiments. All data are mean ± SEM; ***p<0.001 **p<0.01 *p<0.05 relative to WT (unpaired t-test)

Because our data indicated that Pat1 facilitates cell-cell spread, we further investigated whether Pat1 influences actin-based motility, which is known to contribute to spread (63, 64). We found that the *pat1*::Tn mutant formed significantly fewer actin tails compared to WT bacteria at 24 hpi and 48 hpi (**Fig. 6F, G**), suggesting fewer bacteria initiated actin-based motility. Complementation of the *pat1*::Tn (*pat1*::Tn *pat1+*) mutant restored the frequency of actin tail formation to WT levels (**Fig. S3A, B**). In the mixed cell assay, which distinguishes between primary and secondary cells, ∼6% of WT bacteria in the primary cell assembled actin, mostly as actin tails but also as “clouds” of actin surrounding the bacteria, compared with ∼1% of *pat1*::Tn mutant bacteria (**Fig. S3C**). Differences between WT and *pat1*::Tn bacteria in the secondary cell could not be discerned. The observed differences between WT and the *pat1*::Tn mutant were not due to differences in the localization of the *R. parkeri* protein Sca2, which is important for actin-based motility and cell-cell spread (64, 68) (**Fig. S3D**). Taken together, these results suggest that Pat1 is important for the frequency of bacterial actin-based motility, and hence bacterial spread to neighboring cells.

### Pat1 is important for avoiding double membranes during cell-cell spread

Because Pat1 played an important role escaping the primary vacuole following invasion, we hypothesized that it also played a role in escaping the secondary vacuole following cell-cell spread. To test this, we imaged infected HMECs by TEM at 48 hpi and quantified the percent of intracellular bacteria free in the cytosol or within membranes. Significantly more *pat1*::Tn mutant bacteria were surrounded by double membranes (∼60%) in comparison with WT bacteria (∼25%) (**Fig. 7A, B**). The double membranes we observed were often discontinuous, with the mutant remaining mostly enclosed and WT bacteria having very few surrounding membrane fragments. This suggests that Pat1 plays a role in escaping from membranes later in infection.

**Fig. 7.**
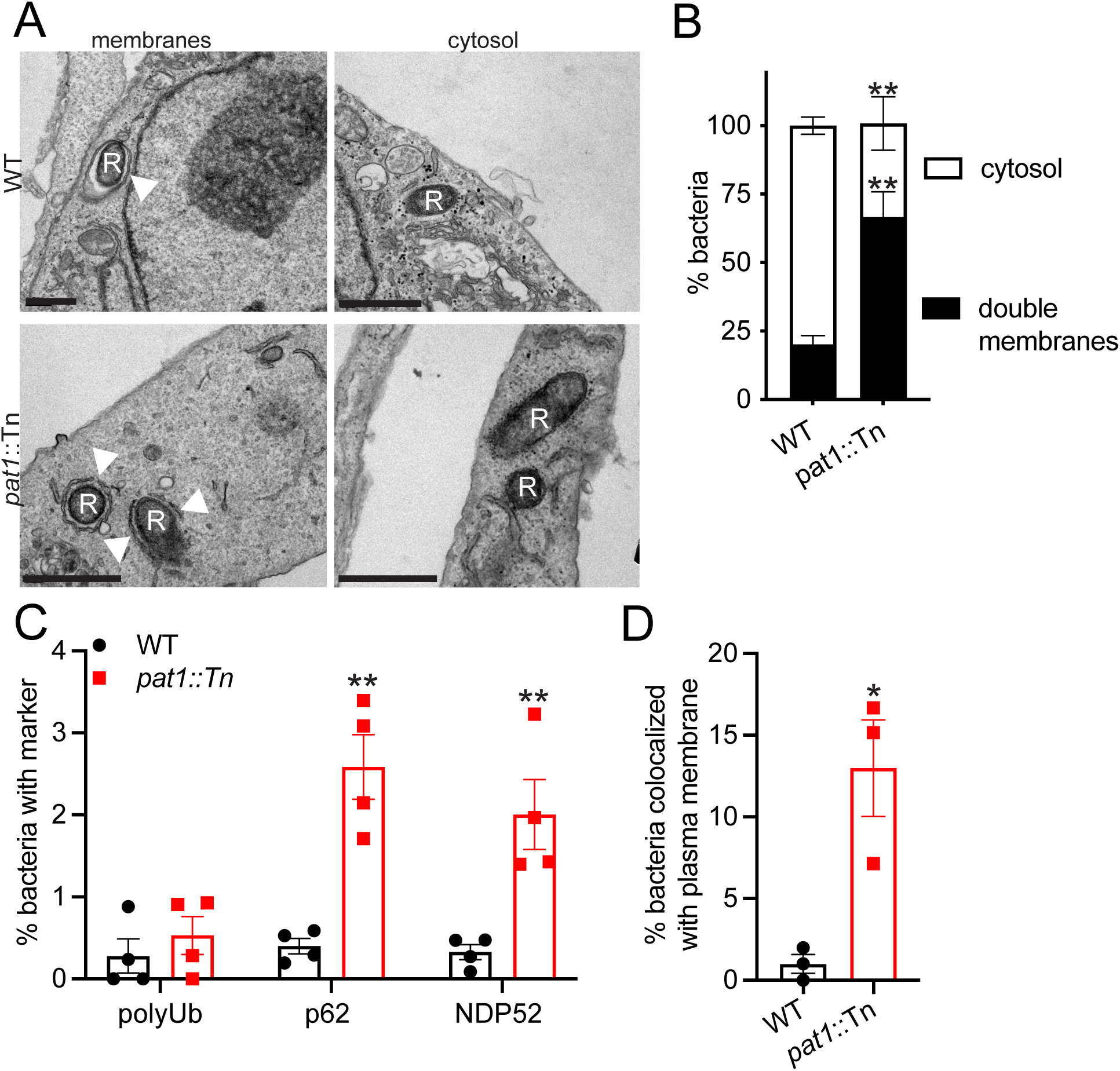
Pat1 is important for escape from the secondary vacuole. (A) TEM images of WT and *pat1*::Tn mutant bacteria in HMECs at 48 hpi. “R” indicates *R. parkeri* and arrowheads point membranes surrounding the bacteria. Scale bar 1 µm. (B) Percentage of bacteria in double membrane compartments or in the cytosol (WT n=120 bacteria, *pat1::tn* n=112 bacteria, n=3 independent experiments). (C) Percentage of bacteria in the secondary cell that colocalize with the plasma membrane from the primary cell, in mixed cell assays from Figure 5C at 32 hpi (n=3 independent experiments). (D) Percentage of WT and *pat1*::Tn mutant bacteria colocalizing with polyUb, p62, and NDP52 (all n=4 independent experiments) at 48 hpi in HMECs. All data are mean ± SEM; **p<0.01 *p<0.05 relative to WT (unpaired t-test).

We next sought to further distinguish whether bacteria surrounded by double membranes were in secondary vacuoles that result from cell-cell spread, or other double-membrane structures involved in autophagy. To determine targeting by autophagy at 48 hpi, we assessed whether bacteria colocalized with polyubiquitin, p62 or NDP52 at 48 hpi. Significantly more of the *pat1*::Tn mutant colocalized with p62 and NDP52 than WT, although the overall percentages were low in all cases (**Fig. 7C**). Moreover, the percentage of bacteria that colocalized with these markers was lower than at 1 hpi (compare with **Fig. 3B, 4C**). Interestingly, polyubiquitin labeling was not significantly different between WT and *pat1*::Tn mutant bacteria, suggesting the observed differences in colocalization with autophagy receptors p62 and NDP52 between WT and the *pat1*::Tn mutant were not mediated by differences in polyubiquitin recruitment. These data suggest that a minor fraction of double membranes observed by TEM at 48 hpi are autophagosomes and Pat1 plays a minor role in autophagy avoidance at late timepoints. To determine whether there instead were differences in escape from secondary vacuole, we used the mixed cell assay described above, in which infected primary cells stably expressing TagRFP-T-farnesyl were infected for 1 h and then mixed with uninfected and unlabeled secondary cells (**Fig. 6C, D**). Fewer than 1% of WT bacteria that spread from primary into secondary cells were colocalized with the plasma membrane marker from the primary cell (**Fig. 7D**), suggesting that these bacteria had escaped the secondary vacuole. In contrast, of the *pat1*::Tn mutant bacteria that spread into secondary cells, ∼12% colocalized with the plasma membrane marker from the primary cell. These results suggest that a significant fraction of double-membrane structures seen in the TEM images are secondary vacuoles and confirm that Pat1 is important for escaping from these vacuoles.

## Discussion

The ability of *Rickettsia* species to escape the vacuole and avoid host membranes is a critical facet of their life cycle. Here, we demonstrate that the *R. parkeri* patatin-like phospholipase, Pat1, is important for virulence in a mouse model of infection. At the cellular level, we find that Pat1 enables bacterial escape from host membranes throughout infection. Pat1 mediates efficient exit from primary vacuoles following invasion, helping *R. parkeri* avoid detection by host galectins and autophagy adaptor NDP52. Pat1 further enables cytosolic bacteria to avoid recruitment of polyubiquitin and autophagy adaptor p62. As infection progresses, Pat1 facilitates spread into neighboring cells and escape from the secondary vacuole. Altogether, these data suggest Pat1 is important at multiple steps of the *Rickettsia* life cycle that involve manipulating host membranes.

Our work shows that Pat1 is important for virulence upon i.v. infection in a mouse model that succumbs to *R. parkeri* infection (8, 61). This is consistent with other bacterial factors involved in vacuolar escape being important virulence factors in animal models of disease. For example, *L. monocytogenes* mutants lacking LLO are avirulent upon i.v. infection in a mouse model and single *L. monocytogenes* PLC mutants are also diminished in virulence (27). Our data suggests that Pat1 plays an important role in *R. parkeri* pathogenesis.

Using TEM and confocal microscopy, we found that Pat1 mediates escape from both single and double membrane compartments in host cells. At early time points, *pat1*::Tn mutant bacteria were more frequently surrounded by single membranes following invasion, likely to be primary vacuoles derived from the host cell plasma membrane. Consistent with a failure to fully escape the primary vacuole, the *pat1*::Tn mutant also showed a reduced frequency of actin-based motility and increased trafficking to LAMP-1-positive compartments. We also found the *pat1*::Tn mutant had increased localization to double membrane structures at later time points when bacteria are spreading to neighboring cells. These structures are likely to be secondary vacuoles, as only a small portion colocalized with autophagy receptors p62 or NDP52. Pat1 was previously suggested as a candidate for escape from the vacuole due to its phospholipase activity (28, 37) and the observation that *R. typhi* pre-treated with anti-Pat1 antibody (which could block surface-associated by not secreted Pat1) caused increased colocalization with LAMP-1 (37). Our results provide genetic confirmation of this role. Several other bacterial phospholipases mediate membrane rupture (69), including *L. monocytogenes* PLCs (19, 26, 27, 69), *Clostridium perfringens* alpha-toxin (PLC) (69–71), and *Psuedomonas aeruginosa* ExoU (69, 72). Similarly, lecithin:cholesterol acyltransferase (LCAT) enzymes from pathogenic protists, including *Plasmodium berghei* phospholipase (PbPL) (73, 74) and *Toxoplasma gondii* TgLCAT (75), have PLA2 and acyl transferase activity that facilitate breakdown of the parasitophorous vacuole. Phospholipases are also used by nonenveloped viruses to breech the endosome (76), including parvovirus capsid protein VP1 which has PLA2 activity (77), and host PLA2 group XVI which is recruited by picornaviruses to endosomes for genome translocation (78). Thus, the role of Pat1 in vacuolar breakdown and escape is similar to functions of phospholipases in other intracellular pathogens.

Despite its importance in escaping from primary and secondary vacuoles, Pat1 is not important for growth in the cell line we tested, suggesting that other *R. parkeri* proteins help rupture vacuolar membranes and gain access to nutrients in the cytosol. Consistent with this notion, the *pat1*::Tn mutant colocalizes more frequently with damaged membranes marked by Gal3 and Gal8. Pat1 must therefore share functional redundancy with other proteins that also function in escape. This is similar to the redundancy between *L. monocytogenes* PLC enzymes, which also have overlapping roles in escape, with double mutants deficient in both PlcA and PlcB showing more severe defects in escape (19, 26, 27) and growth (27, 49) than single mutants. Other *Rickettsia* factor(s) involved in escape may include TlyC, a putative hemolysin (31, 79), which could function analogously to LLO. Pat2, a second PLP, may also have an overlapping role with Pat1 in species like *R. typhi* (28, 37). In addition to the membranolytic proteins, Risk-1, a phosphatidylinositol 3-kinase, was recently reported to manipulate early trafficking events important for invasion, vacuolar escape, and autophagy (80). Factors such as these are likely to function with Pat1 to allow complete escape from vacuolar membranes into the cytosol.

A key role for Pat1 in promoting efficient escape from damaged host membrane remnants is to enable subsequent avoidance of targeting of these membranes by Gal proteins and the autophagy machinery. Consistent with this, we observed that the *pat1*::Tn mutant colocalizes more frequently with Gal3 or Gal8 and the autophagy receptor NDP52. Interestingly, Gal3, but not Gal8, has been found to promote *L. monocytogenes* replication during infection by suppressing autophagy (53). During Group A *Streptococcus* infection, Gal3 has also been shown to prevent recruitment of Gal8 and parkin, which themselves play anti-bacterial roles (81). Whether differential recruitment of Gal3 and Gal8 leads to different outcomes during *Rickettsia* infection remains unknown.

We found that Pat1 plays a role in avoiding targeting by autophagy for bacteria that are not associated with damaged membranes. In support of this notion, the *pat1*::Tn mutant was subject to polyubiquitylation and p62 recruitment without co-recruitment of Gal proteins. Pat1 may thus augment other rickettsial autophagy-avoidance mechanisms, which include lysine methylation of OmpB and OmpB-mediated shielding of bacterial surface from polyubiquitylation (82, 83). In this role, Pat1 might function in a similar manner to PlcA from *L. monocytogenes*, which reduces PI(3)P levels to block autophagosome formation and stall autophagy (49, 50). Both PlcA/B and Pat1 are secreted and can act at a distance, as it was previously shown that a *plcA/B* mutant can be rescued by co-infection with by WT *L. monocytogenes* (50), and we also observed rescue of a *R. parkeri pat1*::Tn mutant by co-infection with WT bacteria. Thus, secreted Pat1 might also target early and/or regulatory aspects of autophagy.

We further found that Pat1 is important for cell-cell spread, including in late actin-based motility and escape from the secondary vacuole (the latter is discussed above). The *pat1*::Tn mutant formed fewer actin tails and exhibited reduced spread into neighboring cells when compared with WT, consistent with the known role for motility in cell-cell spread of SFG *Rickettsia* (63, 64, 68). One key contribution of Pat1 to actin-based motility is to mediate escape from the vacuole, allowing recruitment of the host actin machinery to the surface of the bacteria. However, it remains possible that Pat1 targeting of phosphoinositides (PIs) might also affect actin-based motility, as PIs influence the activity of actin-binding proteins (84–86). Moreover, Pat1 targeting of PIs at the plasma membrane could contribute to protrusion dynamics during cell-cell spread. PIs are implicated in the recruitment and function of the endocytic machinery (87–90). In turn, endocytic pathways have been shown to mediate protrusion engulfment for *L. monocytogenes* and *S. flexneri* (87, 88). Pat1-mediated local membrane damage might also promote spread, as *L. monocytogenes* LLO-mediated membrane damage in the protrusion has been shown to enable exploitation of efferocytosis for spread (89). Thus, Pat1 may play multiple roles in cell-cell spread.

Our data demonstrate that Pat1 plays important roles throughout the *R. parkeri* intracellular life cycle in escaping from membranes, avoiding autophagy, and enabling cell-cell spread. Whether Pat1 also performs other functions during infection remains unknown. For example, secreted Pat1 could contribute to the release of bioactive lipids such as eicosanoids derived from arachidonic acid. Eicosanoid synthesis can impact immunity, inflammation, and vascular function (90–92), and thus represents an underexplored process that may be influenced by *Rickettsia* infection and disease. Membranes are critical hubs of signaling and protein-protein interactions and *R. parkeri*, like other intracellular pathogens, has likely evolved diverse ways of manipulating membranes. Further studies of Pat1 function will elucidate how PLA2 enzymes facilitate microbial adaptation to host cells and could reveal previously unappreciated strategies of membrane manipulation by bacterial pathogens.

## Materials and methods

### Mammalian cell lines

Mammalian cell lines were obtained from the UC Berkeley Cell Culture Facility and grown at 37°C with 5% CO2. Vero cells (African green monkey kidney epithelial cells, RRID:CVCL_0059) were grown in Vero media (DMEM with high glucose (4.5 g/L; Gibco, catalog number 11965-092) and 2% FBS (GemCell, 100500) for culturing or 5% FBS for plaque assays (described below)). A549 cells (human lung epithelial cells, RRID:CVCL_0023) were grown in A549 media (DMEM (Gibco, 11965-092) with high glucose (4.5 g/L) and 10% FBS (ATLAS, F-0500-A)). A549 cells stability expressing a farnesyl tagged TagRFP-T (A549-TRTF) to mark the plasma membrane were described previously (67) and were also maintained in A549 media. HMEC-1 cells (human microvascular endothelial cells RRID:CVCL_0307) were grown in HMEC media (MCDB 131 media (Sigma, M8537) supplemented with 10% FBS (HyClone, SH30088), 10 mM L-glutamine (Sigma, M8537), 10 ng/ml epidermal growth factor (Corning, 354001), 1 ug/mL hydrocortisone (Spectrum Chemical, CO137), and 1.18 mg/mL sodium bicarbonate).

### Plasmid construction

For complementation of *pat1*::Tn *pat1+* mutant, we constructed a pMW1650-Spec-*pat1* complementation plasmid. The plasmid pMW1650-Spec was derived from pMW1650 (93) by deleting the gene encoding GFP and replacing the gene conferring resistance to rifampicin with the *aadA* gene from *E. coli* to confer resistance to spectinomycin (94). Nucleotides 901,999-903,853 from *R. parkeri* genomic DNA were then amplified by PCR and inserted into pMW1650-Spec at the single PstI site. Primers used to amplify this region (5’-ATTGCGACACGTACTCTGCAGATCTCATACCATCATAGTTATAATATTAGC-3’ and 5’-AGAGGATCCCCATGGCTGCAGACACAGGTGTCGTCATTGTGA-3’) contained 15-bp overhang with homology to the plasmid for InFusion (Takara Bio, 638947) cloning and retained the Pst1 site. The amplified sequence contained a predicted promoter 5’ to the *pat1* coding sequence (determined using SoftBerry, BPROM prediction of bacterial promoters; http://www.softberry.com/berry.phtml?topic=bprom&group=programs&subgroup=gfindb) (95) and several predicted transcriptional terminators 3’ to the *pat1* coding sequence (determined using WebGeSTer DB; http://pallab.serc.iisc.ernet.in/gester/) (96).

To express Pat1 in *E. coli* for antibody generation, the DNA sequence encoding full length *pat1* was amplified from *R. parkeri* genomic DNA by PCR and subcloned into a pET1 vector containing an N-terminal 6x His-tag, maltose binding protein (MBP) tag, and TEV cleavage site (Addgene plasmid #29656). Primers used for amplifying the insert (5’-TACTTCCAATCCAATGTAGATATAAACAACAATAAGATTAGC-3’ and 5’- TTATCCACTTCCAATGAGATAACCTTGTACATCATCTGTATGC-3’) contained 15-bp overhang with homology to the plasmid for InFusion cloning (Takara Bio, 638947). The resulting plasmid, pET-M1-6xHis-MBP-TEV-Pat1, was transformed into *E. coli* strain BL21 codon plus RIL-Cam^r^ (DE3) (UC Berkeley QB3 Macrolab).

To express Pat1 in *E. coli* for antibody affinity purification from rabbit sera, the DNA sequence encoding full length *pat1* was amplified from pET-M1-6xHis-MBP-TEV-Pat1 by PCR and subcloned into the pSMT3 plasmid containing a 6xHis-tag upstream of the SUMO tag (97). Primers used for amplifying insert (5’- CACAGAGAACAGATTGGTGGATCCATGGTAGATATAAACAACAATAAGATTAG-3’ and 5’-GTGGTGGTGGTGGTGTAACTCGAGGAGATAACCTTGTACATCATCTGTATGC-3’) contained 15-bp overhang with homology to the plasmid for InFusion cloning (Takara Bio, 638947). The resulting plasmid, pSMT3-6x-His-SUMO-Pat1, was transformed into *E. coli* strain BL21 codon plus RIL-Cam^r^ (DE3) (UC Berkeley QB3 Macrolab).

To make pmCherry-N1-Gal8, full-length Gal8 was amplified from PB-CAG- mRuby3-Gal8-P2A-Zeo (Addgene plasmid #150815) and subcloned into pmCherry-N1 (Clontech, 632523). Primers used for amplifying insert were 5’-ACCGCGGGCCCGGGATCCGCCACCATGATGTTGTCCTTAAACAACC-3’ and 5’-GCGACCGGTGGATCCCCCCAGCTCCTTACTTCCAGT-3’. The forward primer contained a Kozak sequence and both primers contained 15-bp overhang with homology to the plasmid for InFusion cloning (Takara Bio, 638947). To make pmCherry-N1-Gal3, Gal3 cDNA was amplified using primers 5’-CCGGAATTCGCCACCATGGCAGACAATTTTTCGCTC-3’ and 5’-

CGCGGATCCCGTATCATGGTATATGAAGCACTG-3’ and subcloned into pmCherry-N1.

### *R. parkeri* strains and bacterial isolation

*R. parkeri* Portsmouth strain (WT) was provided by Dr. Christopher Paddock (Centers for Disease Control and Prevention). The *pat1*::Tn mutant was generated from this strain as described previously (60). *R. parkeri* strains were purified by infecting confluent Vero cells in T175 flasks at an MOI of 0.05. Flasks were monitored for plaque formation and harvested when 70-80% of the cells in the flask were rounded, typically 5-7 d after infection. Cells were scraped and pelleted at 12,000 x g for 30 min at 4°C. The pelleted cells were resuspended in ice-cold K36 buffer (0.05 M KH_2_PO_4_, 0.05 M K_2_HPO_4_, pH 7.0, 100 mM KCl, 15 mM NaCl) and transferred to a Dounce homogenizer. Repeated douncing of 60-80 strokes released intracellular bacteria and the lysed cells and bacteria were centrifuged at 200 x g for 5 min at 4°C. The supernatant containing the bacteria was overlaid on a 30% MD-76R solution (Bracco Diagnostics, NDC 0270-0860-30) and centrifuged at 18,000 rpm for 30 min at 4°C in a SW-28 rotor to further separate host cel components from bacteria. Bacterial pellets were resuspended in brain heart infusion (BHI) media (BD Difco, 237500) and stored at -80°C. Purified bacteria are referred to as “30% preparations” below.

For complementation of the *pat1*::Tn mutant, small scale electroporations were performed with pMW1650-Spec as previously described for pMW1650 (60). Following electroporation, 200 µl of bacteria per well were added to confluent Vero cells in a 6-well plate. The plate was rocked at 37°C for 30 min in a humidified chamber. An overlay of Vero media with 5% FBS and 0.5% agarose was added to each well and incubated at 33°C, 5% CO_2_ for 24 h. To select for transformants, an overlay of Vero media with 5% FBS, 0.5% agarose, and 40 µM spectinomycin was added to the cells for plaque isolation. Individual plaques were picked and resuspended in 200 µl BHI media. To grow plaque-isolated bacteria, plaques were added to Vero cells in a T25 flask and rocked at 37°C for 30 min. Spectinomycin was added to 40 µM final concentration, and the flasks were placed at 33°C and monitored for bacterial growth and harvested when 70-80% of the cells in the flask were rounded, typically 5-7 d after infection. Infected cells were scraped from the flask and pelleted at 2,000 x g for 5 min at room temperature followed by resuspension in K36 buffer. Cells were mechanically disrupted by vortexing at ∼2,900 rpm (Vortex Genie 2) with 1 mm glass beads for two 30 s pulses with 30 s incubations on ice after each pulse. Following disruption, host cell debris was pelleted by centrifuging at 200 x g for 5 min at 4°C. The supernatant was transferred to pre-chilled microcentrifuge tubes and spun at 10,000 x g for 2 min at 4°C to pellet *R. parkeri*. Bacterial pellets were washed three times with cold 250 mM sucrose, then resuspended in 200 µl BHI (50 µl was frozen at -80°C). To further expand the bacterial population, 150 µl of bead-prepped bacteria was mixed with 350 µl of Vero media and added to Vero cells in a T75 flask, rocked at 37°C for 30 min, supplemented with Vero media to a final volume of 12 ml, and incubated at 33°C in 5% CO_2_. After 5-7 d, when 70-80% of the cells in the flask were rounded, the process of bead disruption and bacteria isolation was repeated, except the bacteria were resuspended in BHI without any sucrose washes to generate frozen stocks that were stored at -80°C. Bacterial strains were screened by PCR for the following: (1) the presence of the original transposon using primers for the rifampicin resistance cassette (primers 5’-ATGGTAAAAGATTGGATTCCTATTTCTC-3’ and 5’-CCTTAATCTTCAATAACATGT-3’); (2) the presence of second transposon using primers for spectinomycin resistance cassette (primers 5’-TGATTTGCTGGTTACGGTGAC-3’ and 5’-CGCTATGTTCTCTTGCTTTTG-3’); and (3) the presence of *pat1*::Tn and WT *pat1* using primers amplifying *pat1* (primers 5’-GTAGATATAAACAACAATAAGATTAGC-3’ and 5’-GAGATAACCTTGTACATCATCTGTATGC-3’). Strains were also screened by assessing plaque size and Pat1 expression by western blot (procedure described below). The *pat1*::Tn *pat1^+^* strain that contained the original transposon, the second transposon, plaque size similar to WT, and restored Pat1 protein levels was propagated to purify bacterial 30% preparations as described above.

The insertion site of the transposon was mapped as previously described (93). Briefly, *R. parkeri* genomic DNA was purified from frozen 30% preparations of bacteria (∼10^8^ PFU). Bacteria were thawed and centrifuged at 8,000 rpm for 5 min to pellet bacteria, then genomic DNA was purified using DNeasy Blood and Tissue kit (Qiagen, 69504), according to the manufacturer’s protocol for Gram-negative bacteria (except that the proteinase K incubation was done overnight). 1 µg of *R. parkeri* genomic DNA was digested with HindIII (New England Biolabs, R0104T). The reaction was heat-inactivated and DNA fragments were self-ligated using T4 DNA ligase (New England Biolabs, M0202T). *E. coli* were transformed with the ligation reaction and plated onto Luria-Bertani (LB) agar plates with 40 µM spectinomycin to select for plasmids containing the pMW1650-Spec transposon and flanking regions of genomic DNA. Plasmid DNA was sequenced (primers 5’-ATCTCGCTTTACCTTGGATTCC-3’ and 5’-CTATACGAAGTTGGGCATAC-3’) to determine the genomic location of the insertion site. The insertion site was then confirmed by PCR from genomic DNA using primers that flank the insertion region (5’-AAAGCGGGAATCCAGTAAATC-3’ and 5’-GGCACAGCAGAAATTACTCTTG-3’).

### Plaque assays and growth curves

To determine the titer of purified bacteria, 200 µl of bacteria diluted 10^-3^-10^-8^ in Vero media were added to Vero cells grown in 6-well plates. Plates were rocked at 37°C for 30 min then overlaid with 3 ml of Vero media with 5% FBS and 0.7% agarose. For imaging plaques, neutral red (0.01% final concentration; Sigma, N6264) in Vero media with 2% FBS and 0.5% agarose was overlaid onto cells 5-7 d post infection. Because of differences in timing of plaque formation for the WT and *pat1*::Tn mutant strains, plaque counts for WT and complemented *pat1*::Tn plaques were done at ∼5 d post infection, and those for *pat1*::Tn plaques were done at ∼7 d post infection. Plaques were counted and imaged 24 h after addition of neutral red.

Growth curves were carried out following infection of HMECs at an MOI of 0.01 in 24-well plates. At each time point, media was aspirated from individual wells, cells were washed twice with sterile deionized water, 1 ml of sterile deionized water was added, and cells were lysed by repeated pipetting. Three serial dilutions of the supernatant from lysed cells in Vero media, totaling 1 ml each, were added in duplicate to confluent Vero cells in 12-well plates. Plates were spun at 300 x g for 5 min at room temperature and incubated at 33°C overnight. The next day, media was aspirated and 2 ml of Vero media with 5% FBS and 0.7% agarose was overlaid in each well. Once plaques were visible, an overlay with neutral red was done as described above. Because of differences in timing of plaque formation noted above, plaque counts for WT and complemented *pat1*::Tn plaques were done at ∼5 d post infection and those for *pat1*::Tn plaques were done at ∼7 d post infection.

### Pat1 expression and antibody generation

For expression of 6xHis-MBP-TEV-Pat1, *E. coli* strain BL21 codon plus RIL-Cam^r^ (DE3) with plasmid pET-M1-6xHis-MBP-TEV-Pat1 was grown in LB with 25 mM glucose an OD_600_ of 0.5 and expression was induced with 1 mM IPTG for 1 h at 37°C. Bacteria were pelleted by spinning at 4,000 rpm for 30 min at 4°C, and the pellet was resuspended in MBP lysis buffer (50 mM Tris-HCl, pH 8.0, 300 mM NaCl, 1 mM EDTA) supplemented with 1 µg/ml each leupeptin (MilliporeSigma, L2884), pepstatin (MilliporeSigma, P5318), and chymostatin (MilliporeSigma, E16), and 1 mM phenylmethylsulfonyl fluoride (PMSF, MilliporeSigma, 52332). Bacteria were flash frozen in liquid nitrogen and stored at -80°C. Bacterial cultures were thawed quickly and kept on ice or at 4°C for the remaining steps. Lysozyme (Sigma, L4919) was added to a final concentration of 1 mg/ml followed by a 15 min incubation on ice. Bacteria were subjected to 8 cycles of sonication at 30% power for 12 s bursts, followed by rest on ice for 30 s. Lysed bacteria were spun at 13,000 rpm for 30 min at 4°C. The supernatant was passed three times over a column of 10 ml of amylose resin (New England Biolabs, E8031L). The column was washed with MBP wash buffer (50 mM Tris-HCl, pH 8.0, 300 mM NaCl) by passing 15 column volumes. Bound protein was eluted by adding 2-3 column volumes of MBP elution buffer (50 mM Tris-HCl, pH 8.0, 300 mM NaCl, 0.5 mM DTT, 10 mM maltose) to the column and collecting 500 µl fractions. Fractions were checked for eluted protein by both Bradford assay and SDS-PAGE, and fractions with the highest concentration of protein and a single band at the expected molecular weight for MBP-Pat1 were pooled and concentrated.

To generate rabbit anti-Pat1 antibodies, 1.7 mg of purified MBP-Pat1 was sent to Pocono Rabbit Farm and Laboratory. Immunization was carried out following their 91-day custom antibody production protocol, then extended for an additional 6 weeks for an additional boost and bleed before final exsanguination.

To affinity purify anti-Pat1 antibodies, *E. coli* strain BL21 codon plus RIL-Cam^r^ (DE3) with plasmid pSMT3-6x-His-SUMO-Pat1 was grown, induced for protein expression, and isolated as described above. Bacterial pellets were resuspended in His lysis buffer (20 mM Tris-HCl, pH 8.0, 300 mM NaCl, 10 mM imidazole) supplemented with protease inhibitors PMSF and LPC at the same concentrations as described above. Bacteria were lysed by sonication and the lysate centrifuged as described above. The supernatant was incubated with 2.0 ml of Ni-NTA resin (Qiagen, 30210) for 1 h at 4°C with rotation and the resin was applied to a column. The column was washed with His wash buffer (20 mM Tris-HCL, pH 8.0, 300 mM NaCl, 30 mM imidazole), and protein was eluted from the column in 500 µl aliquots with 2 column volumes of His elution buffer (50 mM NaH_2_PO_4_, pH 8.0, 300 mM NaCl, 250 mM imidazole). The same protocol was followed to purify 6x-His-SUMO from *E. coli* strain BL21 codon plus RIL-Cam^r^ (DE3) transformed with the parental plasmid pSMT3. Purified 6x-His-SUMO or 6x-His-SUMO-Pat1 were coupled to NHS-activated Sepharose 4 fast flow resin (GE Healthcare, 17-0906-01) in ligand coupling buffer (200 mM NaHCO_3_, pH 8.3, 500 mM NaCl) for 2-4 h at room temperature. To remove anti-SUMO antibodies, the 6x-His-Sumo resin was incubated with 10 ml anti-Pat1 serum diluted in binding buffer (20mM Tris-HCl, pH 7.5) and incubated at 4°C for 2 h with rotation. The flow through was collected and added to the 6x-His-SUMO-Pat1 resin and was incubated at 4°C for 4 h with rotation. Bound antibody was eluted with 100 mM glycine, pH 2.5, into tubes containing 120 µl 1M Tris-HCl, pH 8.8, to neutralize to pH 7.5. Eluted fractions were dialyzed in phosphate-buffered saline (PBS; 137 mM NaCl, 2.7 mM KCl, 10mM Na_2_PO_4_, 1.8 mM KH_2_PO_4_, pH 7.4) with 50% glycerol (pH 8.0) overnight at 4°C, concentrated and stored at -20°C.

### Western blotting

For detection of Pat1 in bacterial cell lysates, 30% purified bacteria were boiled in 1x SDS loading buffer (150 mM Tris pH 6.8, 6% SDS, 0.3% bromophenol blue, 30% glycerol, 15% β-mercaptoethanol) for 10 min, resolved on a 10% SDS-PAGE gel, then transferred to a PVDF membrane (Millipore, IPFL00010). The membrane was blocked overnight at 4°C in TBS-T (20 mM Tris, 150 mM NaCl, pH 8.0, 0.1% Tween 20 (Sigma, P9416) plus 5% dry milk (Apex, 20-241). Affinity purified anti-Pat1 antibody was diluted 1:1,000 in TBS-T plus 5% dry milk and incubated with the membrane overnight at 4°C. Anti-RickA (98) was used as a loading control by diluting serum 1:2,000 in TBS-T plus 5% dry milk and incubating at room temperature for 1 h. Membranes were washed with TBS-T for 5 x 5 min intervals at room temperature. Secondary antibody goat anti-rabbit HRP (Santa Cruz Biotechnology, sc-2004) was diluted 1:3,000 in TBS-T plus 5% dry milk and incubated at RT for 30 min, followed by 5 x 5 min washes with TBS-T. To detect secondary antibodies, ECL HRP substrate kit (Advansta, K-12045) was added to the membrane for 45 s at room temperature and developed using Biomac Light film (Carestream, 178-8207).

### Bacterial infections for imaging

Infections were carried out in 24-well plates unless otherwise noted. For immunofluorescence microscopy, 24-well plates containing 12 mm sterile coverslips were used. HMECs were seeded at 2.5×10^5^ cells/well and infected 36-48 h later. A549 cells were seeded at 1.2×10^5^ cells/well and infected 24 h later. For timepoints from 0-2 hpi, an MOI of 3-5 was used for all cell types, and for 24-48 hpi, an MOI of 0.01-0.05 was used.

For the infectious focus assay, an MOI of 0.001 was used. To infect cells, a 30% preparation of *R. parkeri* was thawed on ice prior to infection and immediately diluted into fresh media on ice. Cell media was aspirated, the well was washed once with PBS (Gibco, 10010049), 0.5 ml of bacteria in media was added per well, and the plate was spun at 300 x g for 5 min at room temperature. Media at 33°C was added following centrifugation and infected cells were incubated at 33°C in 5% CO_2_.

For Gal3 and Gal8 imaging experiments, HMECs were transfected with pmCherry-N1-Gal3 or pmCherry-N1-Gal8 using Lipofectamine LTX (Invitrogen, A12621) and incubated at 37°C overnight. The next morning, wells were washed twice with PBS and replaced with fresh, warm HMEC media. Cells were visually examined to confirm 80-100% confluency and the presence of Gal3 or Gal8 expression. Infections were performed a few hours later.

The mixed cell assay was adapted from a prior study (67). Briefly, A549-TRTF cells and unlabeled A549 cells were seeded into 12-well plates at a density of 3×10^5^ cells/ml and grown overnight. The following day, A549-TRTF cells were infected at an MOI of 5 as described above and incubated at 33°C for 1 h. Both infected A549-TRTF cells and unlabeled A549 cells were detached by adding warm citric saline (135 mM KCl, 15 mM sodium citrate) and incubating for 5 min at 37°C. Cells were gently resuspended by pipetting up and down and recovered in A549 media, then washed twice with A549 media. Cells were resuspended in A549 media with 10 µg/ml gentamycin to kill extracellular bacteria. Infected A549-TRTF and unlabeled cells were mixed at a ratio of 1:120, plated on coverslips in a 24-well plate, and incubated in a humidified secondary container at 33°C until 32 hpi.

Hypotonic shock treatment was adapted from a prior study (65). Briefly, HMECs were infected as stated above and incubated for 5 min at 37°C, at which point media was exchanged with a hypertonic solution (10% PEG-1000, 0.5 M sucrose in PBS) and incubated at 37°C for 10 min. Wells were washed gently once with the hypotonic solution (60% PBS), incubated in hypotonic solution at 37°C for 3 min, then incubated in isotonic HMEC media for 15 min at 37°C.

### Immunofluorescence microscopy

All coverslips were fixed for 10 min in fresh 4% paraformaldehyde (Ted Pella, 18505) at room temperature. Coverslips were washed 3x with PBS pH 7.4 and stored at 4°C until staining. All antibodies were diluted in PBS with 2% BSA (Sigma, A9418). All incubations were done at room temperature unless otherwise noted and all coverslips were mounted in Prolong Gold antifade (Invitrogen, P36930) and sealed with nail polish after drying.

Primary antibodies used to stain *Rickettsia* were rabbit anti-*Rickettsia* I7205 (1:300; (99)); gift from T. Hackstadt), rabbit anti-*Rickettsia* OmpB (1:1,000; (82)), and mouse anti-*Rickettsia* 14-13 (1:400; (99)); gift from T. Hackstadt). Primary antibodies were incubated with coverslips for 30 min. Coverslips were washed 3x with PBS and the following secondary antibodies were added for 30 min, protected from light: goat anti-rabbit Alexa 488 (1:400; Invitrogen, A11008), goat anti-rabbit Alexa 404 (1:150; Invitrogen, A31556), goat anti-mouse Alexa 488 (1:400; Invitrogen, A11001), and goat anti-mouse Alexa 404 (1:150; Invitrogen, A31553).

To quantify colocalization with polyubiquitin and autophagy adapters, cells were permeabilized with 0.5% Triton-×100 and washed three times with PBS. Primary antibodies were added for 30 min to 1 h at the following dilutions: mouse anti-polyubiquitin FK1 (1:250; EMD Millipore, 04-262), guinea pig anti-p62 (1:500; Fitzgerald, 20R-PP001), mouse anti-NDP52 (1:300; Novus Biologicals, H00010241-B01P). For staining with rabbit polyclonal anti-LC3 (1:250; Novus Biologicals, NB100-2220SS) and mouse anti-human Lamp1 (1:25; BD Bioscience, 555801), cells were post-fixed in 100% methanol at room temperature for 5 min. After antibody incubations, coverslips were washed 3x with PBS and the following secondary antibodies were added for 30 min and protected from the light: goat anti-mouse Alexa 568 (1:500; Invitrogen, A11004), goat anti-mouse Alexa 488 (1:400; Invitrogen, A11001), anti-guinea pig Alexa 568 (1:500; Invitrogen, A11075), and anti-guinea pig Alexa 488 (1:400; Invitrogen, A11073). Coverslips were then washed three times with PBS.

To quantify the percent of bacteria with actin tails, cells were permeabilized with 0.5% Triton-×100 for 5 min then washed three times with PBS. *Rickettsia* were stained with either anti-*Rickettsia* 14-13 or anti-*Rickettsia* I7205 as described above. After staining for *Rickettsia,* actin was stained with phalloidin-568 (diluted 1:500 in PBS with 2% BSA; Life Technologies, A12380) for 30 min at room temperature. Coverslips were then washed three times with PBS.

To quantify the percent of bacteria with actin tails in the mixed cell assay, cells were permeabilized with 0.1% Triton-×100 for 5 min then washed three times with PBS. *Rickettsia* was detected with the primary antibody mouse anti-*Rickettsia* 14-13 and the secondary antibody goat anti-mouse Alexa 404 as described above. After staining for *Rickettsia,* actin was stained with phalloidin-488 (diluted 1:400 in PBS with 2% BSA; Life Technologies, P3457) for 30 min at room temperature. Coverslips were then washed three times with PBS.

To quantify the size of infectious foci (64, 67), cells were permeabilized with 0.05% Triton-×100 for 5 min, washed three times with PBS, and blocked with PBS containing 2% BSA for 1 h. Coverslips were incubated with anti-β-catenin (1:200; BD Bioscience, 610153) for 1 h at room temperature, then washed three times with PBS, followed by incubation with goat anti-mouse Alexa-568 (1:500; Invitrogen, A11004) for 30 min protected from the light. Cells were subsequently stained to detect *Rickettsia* with anti-I7205 for 30 min and goat anti-rabbit Alexa 488 as described above. Nuclei were stained with Hoechst (1:10,000; Thermo Scientific, 62249) for 15 min. To quantify, the number of infected cells per focus was counted for 10-15 foci per experiment.

### Transmission electron microscopy

HMEC-1 cells were seeded into 6-well plates (1×10^6^ cells per well) and grown for 36 h. Media was aspirated and 2.5 ml of bacteria in media at an MOI of 5 were added. The plates were spun at 300 x g for 5 min at room temperature, then 2.5 ml of warm HMEC-1 media was added to each well, and the plates were placed at 33°C. Time points were taken by aspirating media, washing the well with PBS, and fixing the cells in fixative (2% paraformaldehyde, 2% glutaraldehyde in 0.05M cacodylate buffer, pH 7.2) for 45 min at room temperature. Cells were scraped and pelleted in microcentrifuge tubes and stored in fresh fixative at 4°C until embedding. Samples were embedded in 2% low melt agarose and placed in 2% glutaraldehyde in 1M cacodylate buffer, pH 7.2, and stored at 4°C overnight. The next day, samples were post-fixed with 1% osmium tetraoxide and 1.6% potassium ferricyanide, then dehydrated in increasing concentrations of ice-cold ethanol (70%-100%). Samples were embedded in Epon 812 resin (11.75g Epon 12, 6.25g dodecenyl succinic anhydride, 7g nadic methyl anhydride, and 0.375 ml of the accelerator benzyldimethylamine was added during the dehydration step) and stained with 2% uranyl acetate and lead citrate. Images were captured with a FEI Tecani 12 transmission electron microscope and analyzed manually to determine the total number of intracellular bacteria and their respective localizations within the cell.

### Mouse Studies

Animal research was conducted under a protocol approved by the University of California, Berkeley Institutional Animal Care and Use Committee (IACUC) in compliance with the Animal Welfare Act and other federal statutes relating to animals and experiments using animals (Welch lab animal use protocol AUP-2016-02-8426-1). Mice were 8-20 weeks old at the time of initial infection. Mice were selected for experiments based on their availability, regardless of sex, and both sexes were used for each experimental group. All mice were of the C57BL/6J background were double knock outs for the genes encoding the receptors for IFN-I (*Ifnar1*) and IFN-*γ* (*Ifngr1*) (*Ifnar1-/-*;*Ifngr1-/-*) (Jackson Labs stock #:029098, described in (61)) and were healthy at the time of infection. *R. parkeri* was prepared by diluting 30% preparation bacteria into 1 ml cold sterile PBS on ice, centrifuging the bacteria at 12,000 x *g* for 1 min, and resuspending in cold sterile PBS to the desired concentration (5 × 10^6^ PFU/ ml for intravenous infection). The bacterial suspensions were kept on ice during injections. Mice were exposed to a heat lamp while in their cages for approximately 5 min and then each mouse was moved to a mouse restrainer (Braintree, TB-150 STD). The tail was sterilized with 70% ethanol, and 200 µl bacterial suspensions were injected using 30.5-gauge needles into the lateral tail vein. Body temperatures were monitored using a rodent rectal thermometer (BrainTree Scientific, RET-3). Mice were euthanized if their body temperature fell below 90°F (32.2°C) or if they exhibited severe lethargy that prevented their normal movement around the cage.

### Statistics

The statistical parameters and significance are reported in the figure legends. Data were considered to be statistically significant when *P* < 0.05, as determined by an unpaired Student’s *t*-test, a one-way ANOVA with either multiple comparisons or comparison to WT bacteria, a two-way ANOVA, or a log-rank (Mantel-Cox) test. Asterisks denote statistical significance as: \**P* < 0.05; \*\**P* < 0.01; \*\*\**P* < 0.001; \*\*\*\**P* < 0.0001, compared with the indicated controls. Statistical analyses were performed using GraphPad PRISM v.9.

## Acknowledgements

We thank Ted Hackstadt, David Wood, and Christopher Paddock for kindly providing strains and reagents. We thank previous Welch Lab members whose work supported this project, including Rebecca Lamason, Natasha Kafai, Julie Choe, and Shawna Reed, and current lab members for critical feedback throughout the development of this project. We also thank the following UC Berkeley core facilities and their facility members for providing equipment, reagents, and technical support to complete this work: Danielle Jorgens, Reena Zalpuri, and Guangwei Min (UC Berkeley Electron Microscope Laboratory); Holly Aaron and Feather Ives (CRL Molecular Imaging Center); and Alison Killilea (Cell Culture Facility). We thank David Drubin, Karsten Gronert, and Daniel Portnoy for technical discussion and critical guidance for this work. We also thank Neil Fischer for proofreading the manuscript. This work was funded by grant R01 AI109044 from the NIH/NIAID to M.D.W.

