## SupplementaryFigures for "A patatin-like phospholipase mediates *Rickettsia parkeri* escape from host membranes"

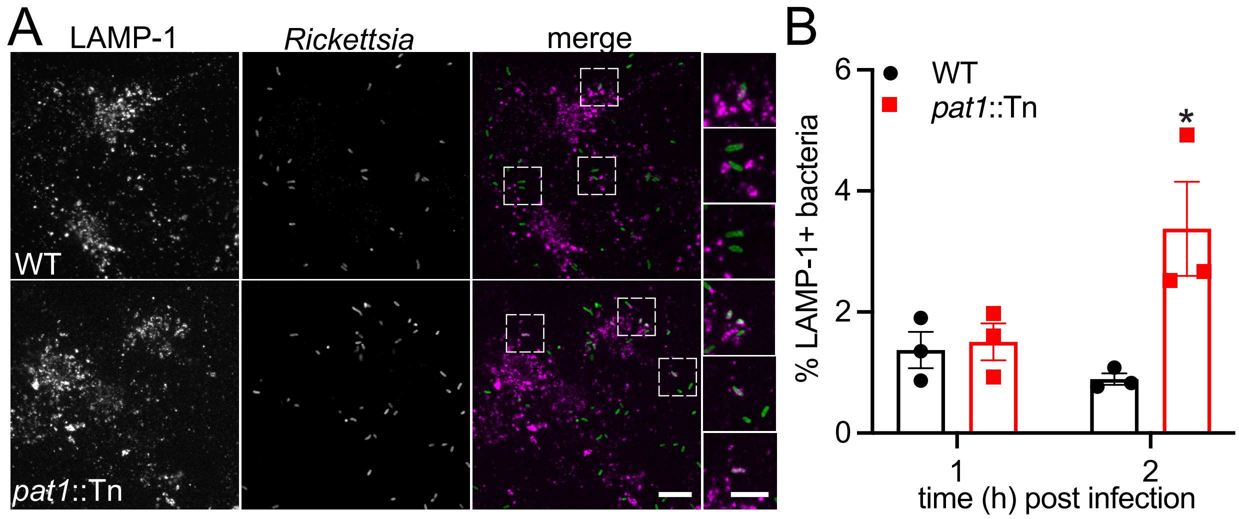


**Fig. S1 Pat1 enables avoidance of trafficking to late endosomal and lysosomal compartments.** (A) Images of LAMP-1 (magenta in merge) in HMECs infected with WT or *pat1*::Tn bacteria (green in merge) at 2 hpi. Boxes indicate insets on right. Scale bar is 10 μm, inset 3 μm. (B) Quantification of LAMP-1 positive bacteria at 1 hpi (images not shown) and 2 hpi (images in (A)). All data represents n=3 independent experiments. Data in (B) are mean ± SEM; *p<0.05 relative to WT (unpaired t-test).

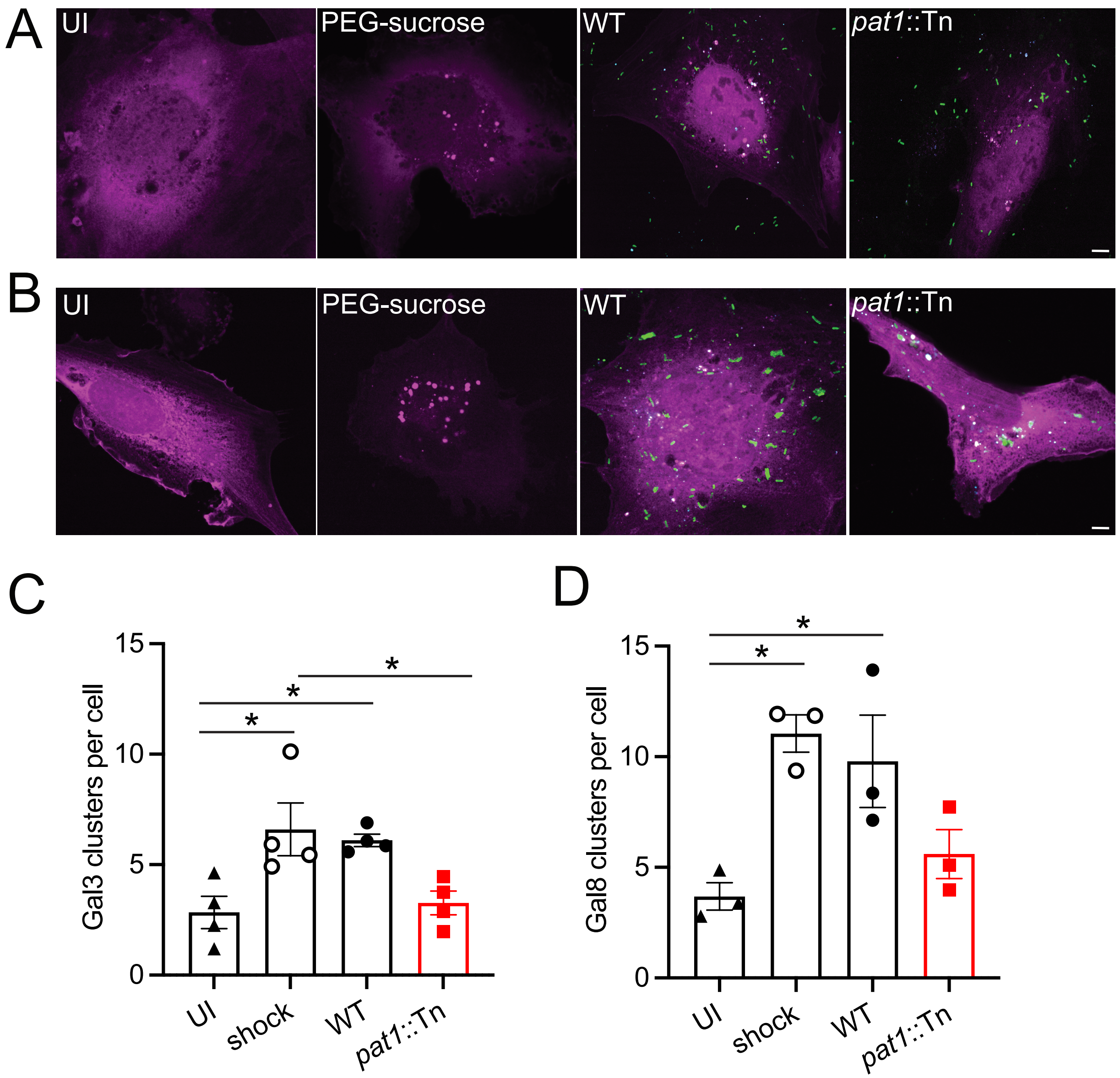


**Fig. S2 Pat1 contributes to intracellular membrane damage.** Images of (A) Gal3-mCherry (magenta) and (B) Gal8-mCherry (magenta) in HMECs that are uninfected (UI), undergo sterile lysis of vesicles (hypotonic shock; PEG-sucrose), WT-infected, and *pat1*::Tn mutant-infected (green) infected at 1 hpi. Infected panels are also stained for NDP52 (cyan). Scale bars for (A) and (B) are 5 μm. (C) Number of Gal3 clusters per cell (n=4 independent experiments). (D) Number of Gal8 clusters per cell (n=3 independent experiments). Data in (C) and (D) are mean ± SEM; *p>0.05 (one-way ANOVA, multiple comparisons with Tukey post hoc test).

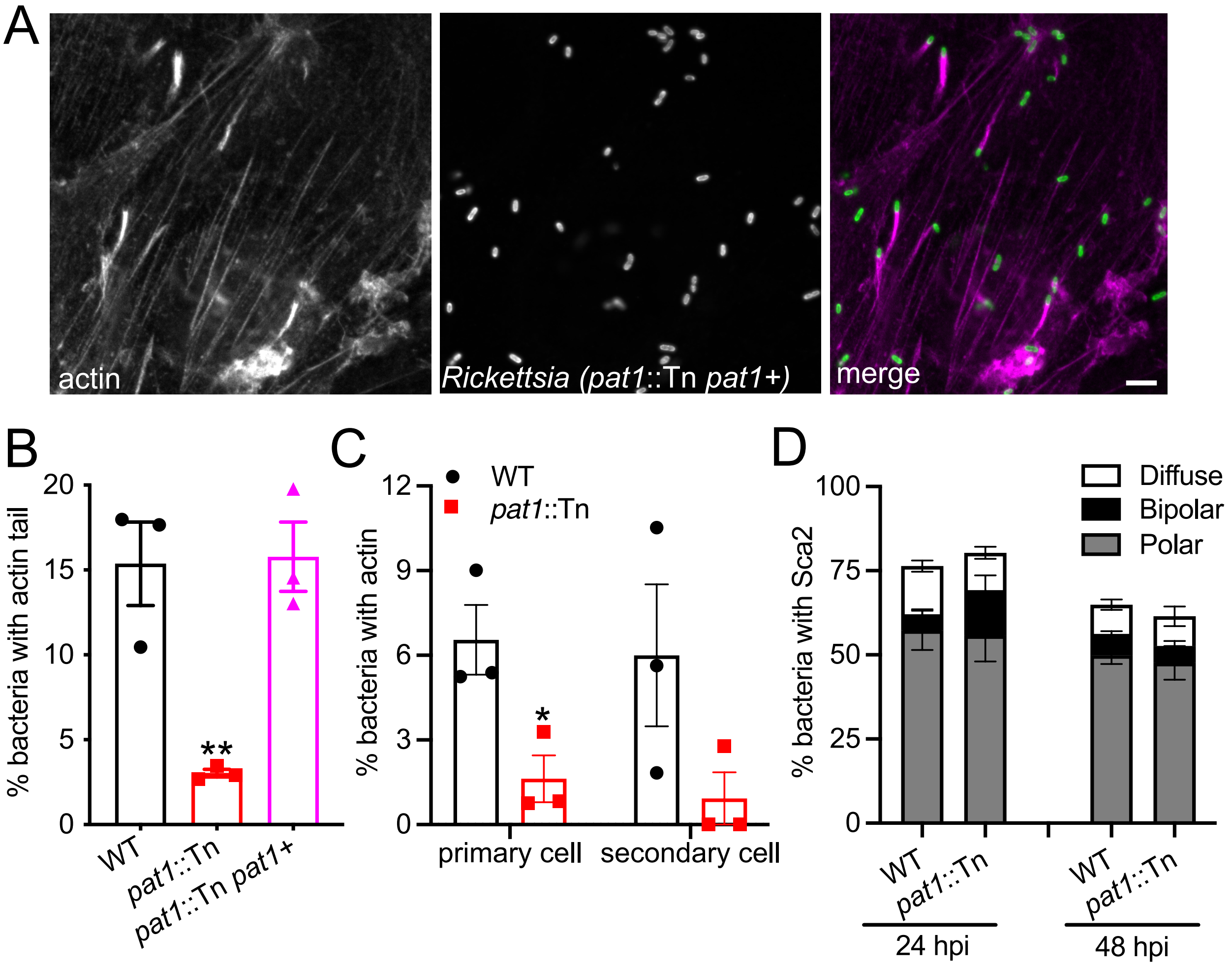


**Fig. S3 The reduced frequency of actin-based motility for the *pat1*::Tn mutant occurs in the primary cell and is not due to altered localization of Sca2.** (A) Images of actin tails in the complemented mutant (F-actin, magenta in merge; bacteria, green in merge) in HMECs at 48 hpi. (WT and *pat1*::Tn images represented in figure 6F.) Scale bar is 5 μm. (B) Percentage of bacteria with actin tails for the indicated strains. (C) Percentage of bacteria with actin tails in primary and secondary cells, related to Figure 5C. (D) Percentage of bacteria with Sca2 with the indicated distributions in WT and *pat1*::Tn mutant bacteria (images not shown). All data represents n=3 independent experiments. Data in (B) are mean ± SEM; **$p<0.01$ relative to WT (one-way ANOVA). Data in (C) are mean ± SEM; *$p<0.05$ relative to WT (unpaired t-test).
